# Epidermal Stem Cells Control Periderm Injury Repair via Matrix-Driven Specialization of Intercellular Junctions

**DOI:** 10.1101/2025.07.02.662640

**Authors:** Helen Mengze He, Liana C. Boraas, Jon M. Bell, Xiangyu Gong, Sophia L. Iannaccone, Zhang Wen, Michael Mak, Marina Carlson, Kaelyn Sumigray, Stefania Nicoli

## Abstract

Epidermal stem cells interact with the extracellular matrix (ECM) to regulate their differentiation and maintain skin architecture. Here, we demonstrate a novel role for basal epidermal stem cells (BECs)-ECM interaction in regulating adhesion molecules expressed by the periderm—the superficial epidermal cells (SECs) of the embryonic bilayered skin. Using the developing zebrafish fin fold, we identify BECs form distinct regions of collagen-versus laminin-enriched basement membranes through integrin-mediated adhesions. Mechanistically, collagen-associated BECs form desmosomes and adherens junctions (AJs) with SECs while laminin-associated BECs display reduced desmosomes but sustain AJs and actomyosin expression with SECs. Notably, we show both *in vivo* and in a bilayered human keratinocyte model, that laminin, compared to collagen, is sufficient to repress desmosome formation while sustaining AJs specifically at the interlayer cell contacts. *In vivo*, laminin deficiency enhances desmosome expression across layers and impairs the wound-healing capacity of SECs. This defect was partially rescued by genetic reduction of the desmosome protein Desmoplakin-1a, highlighting the role of ECM-dependent junctional specialization in mediating differences in SEC injury response.

Overall, our findings identify that stem cells, through their matrix, establish specialized junctions in the overlying stratified epithelium, which contribute to skin healing properties.

## Introduction

Stem cells are essential for tissue development, maintenance, and repair due to their capacity for self-renewal and differentiation^1–3^. Their behavior is regulated by cues from the surrounding microenvironment, or niche, which includes neighboring cells and the extracellular matrix (ECM). They interact with specific ECM components via integrins^4,5^, and form junctions with neighboring cells. For example, adherens junctions (AJs) facilitate intercellular adhesion by connecting the actin cytoskeletons of adjacent cells and modulate tissue tension through actomyosin-driven mechanotransduction^6^. Additionally, desmosomes provide tissue resistance to tearing or separation by firmly linking adjacent cells through intermediate filaments^7^. These junctions work synergistically, and altering any of these interactions can affect stem cell proliferation and differentiation^8–11^. While the roles of these interactions in stem cell differentiation are well established, it remains unclear whether they also allow stem cells to influence the behavior of tissues independent of fate specification. Exploring this aspect could uncover a critical, underappreciated structural role of stem cells in shaping tissue responses.

Basal epidermal stem cells (BECs) originate from the ectoderm during embryonic development and expand over an ECM that they produce, covering the embryo’s surface as early as four weeks post-gestation in humans^12^. BECs differentiate to form a stratified epidermis by approximately eight weeks of gestation. However, there is a critical period during human embryonic development, before week eight ^13^, when BECs do not undergo differentiation or asymmetric division. Instead, they expand and interact with the periderm, a temporary layer consisting of superficial epidermal cells (SECs) that protects the embryo from the amniotic environment^14^. Because BECs are undifferentiated, adhesive, and form stable contacts within a bilayered epidermis, they present a unique model for studying how basal cells contribute to tissue architecture and interlayer organization prior to stratification.

Analyzing and manipulating BECs within their intact environment *in utero* in early mammalian embryos is technically challenging, leaving these functions largely unexplored. Other vertebrate embryos, such as zebrafish, also possess a conserved bilayer epidermis^15–17^. In zebrafish, the fin fold epidermis is bilayered from as early as 2 days post-fertilization (dpf)^17,18^, and is composed of BECs and SECs, which functionally resemble the mammalian periderm.

Notably, the zebrafish fin fold covers the embryonic tailbud and developing limb-like structures, analogous to the periderm coverage of mammalian limb buds^19^. The BECs remain undifferentiated until around 15 dpf, when they begin to differentiate into a stratified epidermis^16^. Thus, we used the zebrafish fin fold, and an *in vitro* human bilayer skin model, to investigate how BECs regulate epidermal architecture and function independently of differentiation.

We discovered that BECs generate distinct ECM zones within the fin fold, characterized by collagen-rich ECM at the center and laminin-rich basement membranes at the periphery. In both zebrafish and human skin models, a laminin-rich ECM beneath BECs was both necessary and sufficient to suppress desmosome formation while maintaining AJs at the BEC–SEC interface. *In vivo*, this junctional specialization promotes tissue repair after cell injury in the wild-type periderm. While impaired collagen decreases both AJs and desmosomes, reduced laminin expression increases desmosome formation in BECs and across the epidermal bilayer, suppresses actomyosin expression at AJs, and impairs wound healing of the periderm at the fin fold periphery. Partial depletion of desmoplakin 1a (*dspa*), which encodes for the desmosomal linker protein expressed across epidermal layers, partially rescues the wound closure defects observed at the fin fold periphery of laminin mutants.

Our work suggests that BECs strengthen periderm resilience to tissue injury by forming region-specific ECM zones that direct intercellular junction specialization, revealing a role in skin organization and repair independent of differentiation.

## Results

### Basal epidermal stem cells (BECs) form distinct zones of extracellular matrices in zebrafish embryos

During zebrafish embryogenesis, the entire embryo is enveloped by a bilayer epidermis (Fig. 1A)^17,20^. The bottom layer consists of basal epidermal stem cells (BECs), which express p63^21–23^, and the top layer of superficial epidermal cells (SECs), called periderm, are p63-negative and express higher levels of keratin 5^21^ (Fig. 1A-C and Extended Data Fig. 1A).

**Figure 1.**
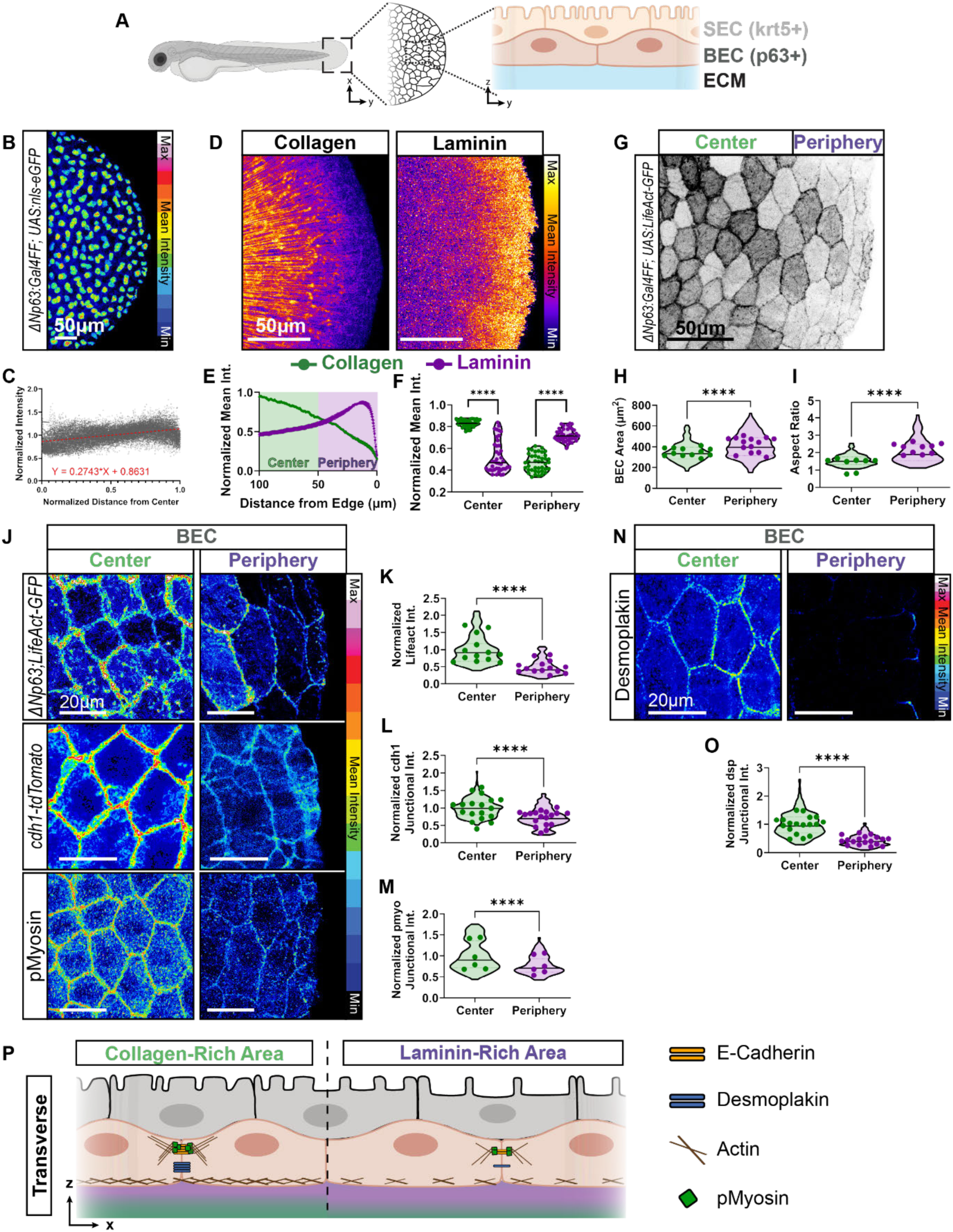
Region-Specific Junctional and Cytoskeletal Organization of BECs in the Fin Fold. (**A**) Schematic of zebrafish fin fold at 2 to 2.5 days post fertilization (dpf). Magnified transverse view showing basal epidermal stem cells (BECs) on basement membrane extracellular matrix (ECM) and peridermal superficial epidermal cells (SECs). (**B**) Representative images of *TgBAC(ΔNP63:Gal4ff)^la2^*^13^*; Tg(UAS:NLS-GFP)^el6^*^09^ for BECs at 48 hours post fertilization (hpf). Intensity displayed using 16-color scale. (**C**) Quantification of nuclear GFP mean intensity across the fin fold normalized to each embryo’s average intensity, plotted against normalized distance from fin fold center (1.0 = edge of the fin fold). Simple linear regression (red dashed line) equation reports slope of 0.2743 (n = 42 embryos, two independent experiments). (**D**) Representative image of FAM-tagged collagen hybridizing peptide (left) and Laminin A1 protein (right). Intensity is displayed using a fire LUT. (**E**) Quantification of normalized intensity from edge of the fin fold (edge = 0 µm). Periphery < 50 µm of the edge, Center> 50 µm. Violin plot of averaged values for center and periphery per embryo shown in (**F**) (Kruskal-Wallis test with multiple comparisons, p<0.0001, n = 41 embryos; 4 independent experiments). (**G**) Representative image of *TgBAC(ΔNP63:Gal4ff)^la2^*^13^*; Tg(UAS:LifeAct-GFP)^mu2^*^71^ in central and peripheral regions as defined in panels E and F. Quantification of (**H)** BEC area and (**I)** aspect ratio displayed as a violin plot representing the distribution of individual cells, with average values per embryo (dots) in both the central and peripheral regions (two-tailed Mann-Whitney test on violin plots, p<0.0001, n=114 - 152 cells, dots n = 8-14 embryos; 2 independent experiments). (**J, N)** Representative images showing (**J**) *TgBAC(ΔNP63:Gal4ff)^la2^*^13^*; Tg(UAS:LifeAct-GFP)^mu2^*^71^ for endogenous labeling of E-cadherin, alongside immunofluorescence staining for phosphorylated myosin light chain and (**N**) desmoplakin1/2 in the central and peripheral regions of fin fold BECs at 48 hpf. Z-stack images obtained from the SEC layer to the BEC layer. BECs identified by maximum projection of selected z slices based on their z-position. Intensity is displayed using a 16-color scale. (**K-M, O**) Quantification of (**K**) BEC-LifeAct-GFP per cell, (**L**) junctional E-cadherin, (**M**) junctional phospho-myosin light chain, and (**O**) junctional desmoplakin1/2 across embryos, with individual distributions shown as violin plots and average values per embryo as dots. Intensity was normalized to the Center group for each graph. (two-tailed Mann-Whitney test on violin plots, p < 0.0001, n=172-175 cells (BEC-LifeAct), 240 junctions (E-cadherin), 93-120 junctions (pMyosin), and 284-348 junctions (Desmoplakin), dots n =13-14 (BEC-LifeAct; 2 independent experiments), 18 (E-cadherin; 4 independent experiments), 6 (pMyosin; 1 independent experiment), and 18 (desmoplakin; 2 independent experiments) embryos). (**P**) Summary schematic of intercellular junctions in basal epidermal stem cells above laminin-vs. collagen-positive basement membranes.

To investigate how undifferentiated BECs contribute to epidermal organization, we first analyzed the composition and spatial distribution of ECM components in the zebrafish fin fold. BECs in the fin express specific types of collagen and laminin^24^, protein components of the extracellular matrix (ECM) that in turn are required for fin fold development^21,25,26^ (Extended Data Fig. 1B-E). Collagen is produced by both BECs and embryonic fibroblasts, in contrast, laminins are primarily produced by BECs (Extended Data Fig. 1F)^27,28^. Confocal imaging of fluorescein-tagged collagen hybridizing peptide, to detect all types of collagens^29^, revealed fibrillar collagens in the stromal ECM in the fin fold central region, which coincides with the localization of fin mesenchymal cells and interstitial fibroblasts in the dermis^30^ (Fig. 1D-F, and Extended Data Fig. 1G, H). We also detected cytoplasmic collagens in BECs in the central region (Extended Data Fig. 1H). Unexpectedly, we found that BECs produce a relatively high level of laminin^31^ only in the peripheral zone of the fin fold compared to the central region (Fig. 1D-F, and Extended Data Fig. 1I).

These findings reveal a region-specific pattern of ECM composition partially produced by BECs depending on their spatial context within the fin fold.

### Peripheral BECs on laminin-rich extracellular matrices have reduced adherens junctions and desmosomes

We hypothesized that regional ECM differences in the fin fold are associated with changes in the cytoskeletal and junctional organization of BECs. To test this, we first analyzed the actin cytoskeleton in BECs. Briefly, we performed confocal live imaging of the actin reporter LifeAct-GFP regulated by the BEC-specific promoter *p63*. Notably, BEC morphologies and areas were significantly different in the central and peripheral regions of the fin fold (Fig. 1G-I). Specifically, BECs above the peripheral, laminin-rich areas exhibited a larger and more elongated cell area and lower actin levels compared to those in the central, collagen-dense regions (Fig. 1H-K).

Actin filaments interact with adherens junctions (AJs), intercellular junctions that regulate tissue tension via phosphorylated myosin^32^. By live confocal microscopy, we found that the expression of the AJ component E-cadherin^33^ was lower in peripheral BECs compared to the central BECs (Fig. 1J, L). Similarly, immunostaining followed by confocal microscopy revealed reduced junctional expression of phosphorylated-myosin and vinculin^34,35^ in peripheral compared to central BECs (Fig. 1J, M and Extended Data Fig. 1J, K). To determine if these differences in BECs were unique to AJs, we examined desmosomes. Immunostaining of desmoplakin 1/2^36^ revealed substantially reduced levels of this desmosome component in the peripheral BECs compared to the neighboring central BECs (Fig. 1N, O).

Overall, these data suggest that BECs have differential adhesion profiles correlated with a region-specific pattern of ECM composition (Fig. 1P).

### Peripheral SECs above laminin-rich ECM have reduced desmosomes but retain high levels of adherens junctions

In zebrafish, as in mammals, BECs are tightly connected to SECs^10,22,37^. Thus, we examined how regional ECM differences beneath the BEC layer might influence the SECs above. Specifically, we analyzed cytoskeletal and junctional organization in SECs overlying the central (collagen-rich) versus peripheral (laminin-rich) zones of the fin fold.

First, we performed confocal live imaging of LifeAct-GFP regulated by the SEC-specific promoter *krt5* (Fig. 2A and Extended Data Fig. 1A). Similar to BECs, SECs exhibited differential cytoskeletal organization across the ECM zones. Specifically, in the central, collagen-rich region, SECs exhibited junctional actin and well-organized microridges, which are apical surface protrusions composed of polymerized actin. In contrast, SECs in the peripheral, laminin-positive region showed similar junctional actin but less organized apical actin (Fig. 2A).

**Figure 2.**
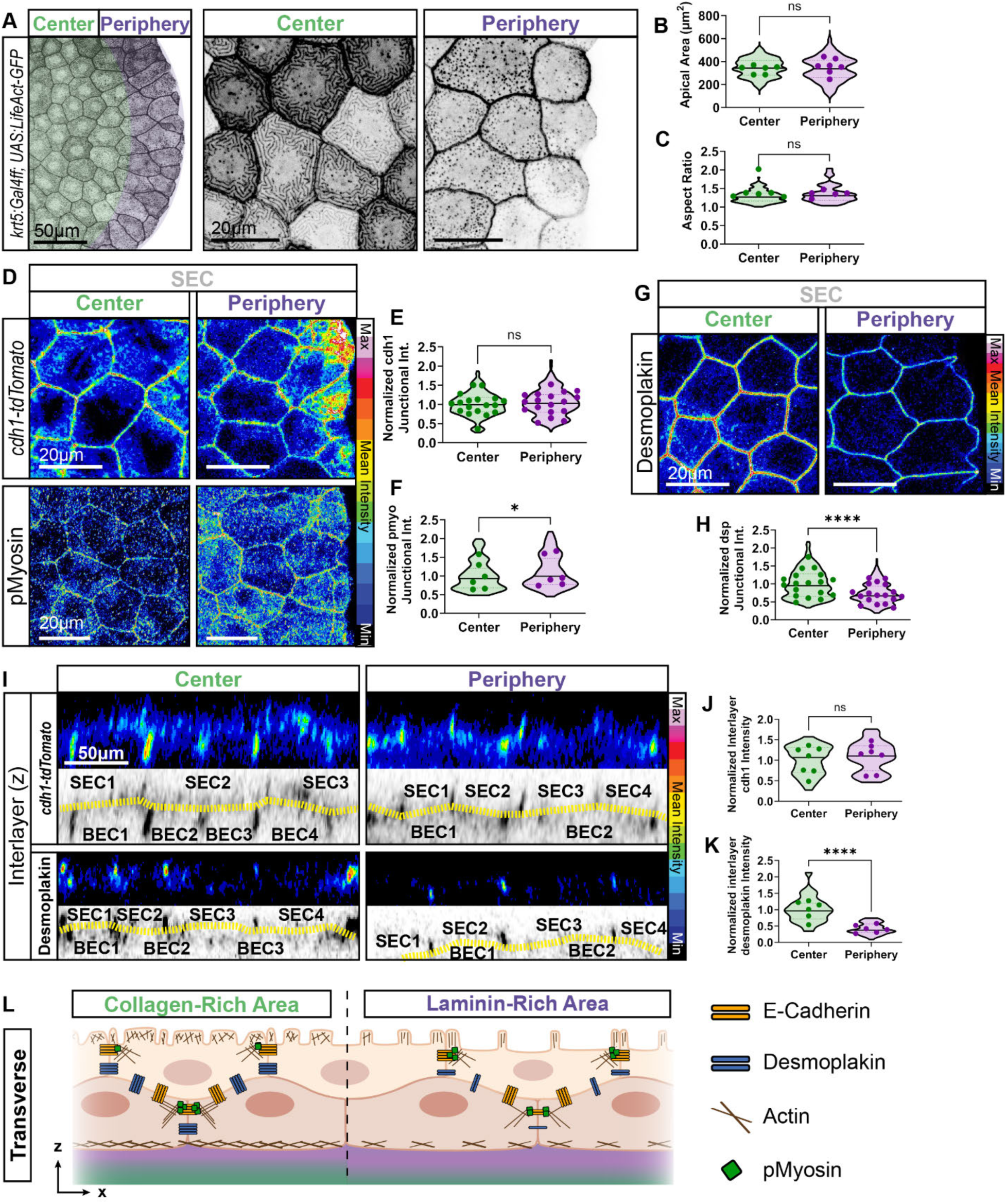
Differential Cytoskeletal and Junctional Organization in SECs Across Central and Peripheral ECM Zones. (**A**) Representative image of *Tg(krt5:Gal4ff)^la2^*^12^*; Tg(UAS:LifeAct-GFP)^mu2^*^71^ highlighting the central region (green) and peripheral boundary (purple) as defined in figure 1D-F. Quantifications of (**B**) SEC apical area and (**C**) aspect ratio with individual cell distribution shown as violin plots and average values per embryo as dots (two-tailed unpaired t test (apical area) and two-tailed Mann-Whitney test (aspect ratio), p = 0.4548 (apical area), 0.1672 (aspect ratio), n =83-86 cells (apical area), 48-160 cells (aspect ratio), dots n = 6-7 (apical area), and 5-6 (aspect ratio), embryos; 3 independent experiments). (**D, G)** Representative images showing (**D**) *cdh1-tdTomato^xt^*^18^, alongside immunofluorescence staining for phosphorylated myosin light chain and (**G**) desmoplakin1/2 at 48 hpf. Z-stack images taken from the SEC layer to the BEC layer. SECs were identified by maximum projection of selected z slices based on their z-position. Intensity displayed using 16-color scale. (**E-F, H**) Quantification of intensity of (**E**) junctional E-cadherin, (**F**) junctional phospho-myosin light chain, and (**H**) junctional desmoplakin1/2 across embryos, individual junctional measurements distribution shown as violin plots and average values per embryo as dots. Intensity was normalized to the Center group for each graph (two-tailed Mann-Whitney test on violin plots, p = 0.2577 (E-cadherin), 0.0457 (pMyosin), and p < 0.0001 (Desmoplakin), n= 372 junctions (E-cadherin), 99-119 junctions (pMyosin), and 362-366 junctions (Desmoplakin), dots n = 18 (E-cadherin; 4 independent experiments), 6 (pMyosin; 1 independent experiment), and 18 (desmoplakin; 2 independent experiments) embryos). **(I)** Representative orthogonal view of the fin fold, showing live imaging of *cdh1-tdTomato^xt^*^18^ or desmoplakin1/2 immunofluorescence staining in collagen-enriched (center) and laminin-enriched (periphery) regions. (Top) Images displayed using 16-color scale. (Bottom) Interface membrane (yellow dashed lines) between SEC and BEC layers, with numbers corresponding to each analyzed cell. **(J, K**) Quantification of interlayer (yellow dashed lines) (**J**) E-cadherin and (**K**) desmoplakin across embryos, individual measurements distribution shown as violin plots and average values per embryo as dots. Intensity was normalized to the Center group for each graph (two-tailed Mann-Whitney test on violin plots, p= 0.2917 (E-cadherin) and p<0.0001 (Desmoplakin), n= 70 (E-cadherin), and 60 cells (desmoplakin), dots n= 7 (E-cadherin; 2 independent experiments) and 6 (desmoplakin; 1 independent experiment) embryos). (**L**) Summary schematic of junctional characteristics in SECs, SEC-BEC interface, and BECs.

Interestingly, SECs in the peripheral and central regions had similar apical cell areas and cell shapes, indicating that the differential distribution of microridges is not associated with cortical contractions^38^ (Fig. 2B, C). Furthermore, SECs in both regions had equal E-cadherin, and phosphorylated myosin at the junction was elevated in peripheral versus central SECs (Fig. 2D-F). In contrast, desmoplakin junctional expression was reduced in peripheral versus central SECs (Fig. 2G-H). These findings suggest that SECs above laminin-rich ECM maintain epithelial integrity through enhanced actomyosin signaling, potentially compensating for reduced desmosomal adhesion. Thus, SECs exhibit region-specific junctional complexity, characterized by a decoupling of AJs and desmosomes (Fig. 2L).

Since intercellular SEC-SEC and BEC-BEC junctions differ in the peripheral and central regions, we investigated whether these molecular differences extended to interlayer adhesion between BECs and SECs. We performed live imaging of the *cdh1-tdTomato; krt5:Gal4ff, UAS:LifeAct-GFP* transgenic line, which allowed us to discern the SEC layer (pseudo-colored in grey and yellow) from the BEC layer (pseudo-colored in yellow only) (Extended Data Fig. 2A, Supplementary Video 1). Notably, the BEC and SEC layers migrated synchronously over the 8 hours of imaging, both at the center and periphery regions, with no observable difference in the mean squared displacement between BECs and SECs (Extended Data Fig. 2B). Further tracking of individual BECs and SECs showed sustained interlayer contact over time (Extended Data Fig. 2C), suggesting their fixed connection in both center and periphery. However, when we examined the levels of junction proteins at the membrane interface between BECs and SECs, we found that desmoplakin was more abundant at the central versus peripheral BEC-SEC interface, and E-cadherin levels were consistent across the entire interface (Fig. 2I-K), similar to the organization of SEC-SEC junctions. These findings suggest that the BEC-SEC bilayer is connected via both desmosomes and AJs above the collagen-rich central region, and mainly via AJs above the laminin-rich peripheral region (Fig. 2L).

Together, these findings support a model in which BEC–ECM interactions direct the assembly of regionally specialized junctions across the bilayer, with peripheral SECs relying more heavily on actomyosin contractility and AJs to maintain tissue cohesion in the context of reduced desmosomal support.

### Integrin-dependent BEC-ECM adhesion promotes adherens junctions and suppresses desmosomes throughout the bilayer epidermis

We hypothesized that BEC-ECM adhesions regulate the organization of intercellular junctions within and between both layers of the epidermis. To test this, we examined mutants with reduced levels of integrin β1, a transmembrane receptor required for cell-ECM adhesions in both collagen- and laminin-rich ECMs^39^.

In zebrafish, integrin β1 is encoded by *itgb1b* and *itgb1a,* and we detected high levels of *itgb1b* mRNA and protein in BECs in both regions, but especially in the central BECs (Extended Data Fig. 3A-E). As expected, we did not detect high levels of *itgb1b* and *itgb1a* in SECs, which do not directly interact with the ECM (Extended Data Fig. 3A-E). To avoid possible compensatory effects between isoforms, we analyzed double heterozygous *itgb1b*;*itgb1a* mutants (*itgb1b^+/-^*;*itgb1a^+/-^)*. These mutant embryos did not exhibit any obvious defects in growth or fin fold development (Extended Data Fig. 3F).

Given that integrins can alter the expression of ECM components ^40,41^, we verified that *itgb1b^+/-^;itgb1a^+/-^*mutant embryos show a fin fold morphology, region-specific ECM composition, and overall ECM levels comparable to wild-type (*WT*) controls (Fig. 3A-C and Extended Data Fig. 3F). This allowed us to isolate the role of integrin-mediated adhesion independent of changes in ECM.

**Figure 3.**
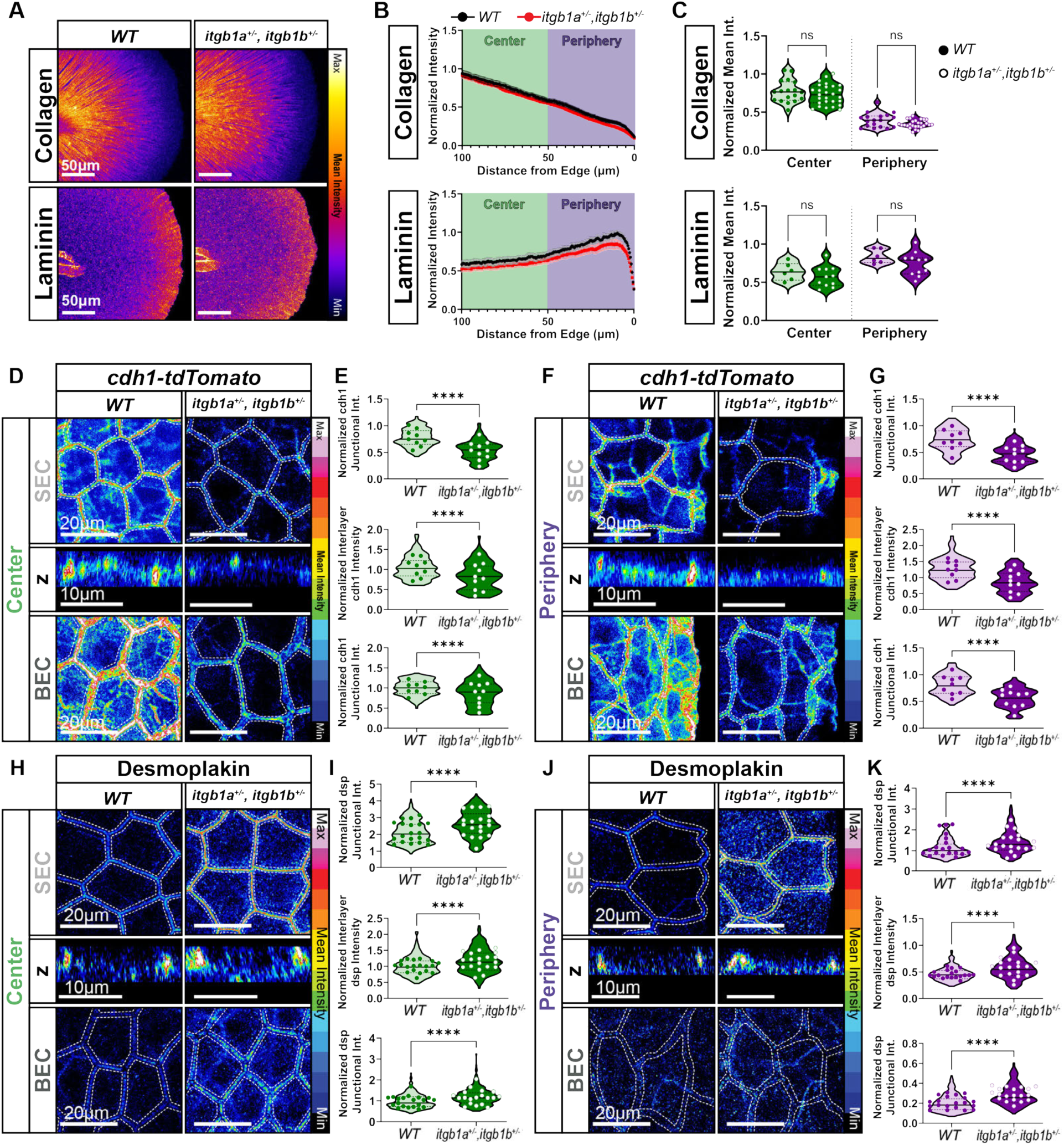
BEC-ECM Adhesions Regulate the Balance Between Desmosome and Cadherin Junctions Across Layers. (**A**) Representative image of of *wild type* (*WT*) vs. *itgb1a^+/-^, itgb1b^+/-^* fin fold, stained with a FAM-tagged collagen hybridizing peptide and Laminin A1 at ∼48 hpf. Intensity displayed using fire LUT. (**B)** Quantification of normalized collagen and laminin intensity across fin fold (edge = 0 µm). Each curve is normalized to individual embryo maximum. (**C**) Quantification of average normalized collagen and laminin intensity per embryo (Collagen: Ordinary One-Way ANOVA test with multiple comparisons, p = 0.6422 (Center) and 0.4573 (Periphery), n = 18 (*WT*) and 27 (*itgb1a^+/-^, itgb1b^+/-^)* embryos; 3 independent experiments; Laminin: Kruskal-Wallis test with multiple comparisons, p>0.9999, n = 6 (*WT*) and 11 (*itgb1a^+/-^, itgb1b^+/-^)* embryos; 2 independent experiments. (**D, F, H, J**) Representative images of *cdh1-tdTomato^xt^*^18^ ((**D**) center and (**F**) periphery) and immunofluorescence staining for desmoplakin ((**H**) center and (**J**) periphery) in zebrafish fin folds at 48 hpf. Images show SECs, Interlayer (between SEC and BEC), and BECs. Z-stack images were taken from the SEC layer to the BEC layer. SECs and BECs were identified by maximum projection of selected z slices based on their z-position. Representative interlayer images are shown in orthogonal view of single z slice. Intensity displayed using 16-color scale. (**E, G, I, K**) Quantification of E-cadherin ((**E**) center and (**G**) periphery) and desmoplakin ((**I**) center and (**K**) periphery) at SEC junctions, interlayer, and BEC junctions with individual junctional measurements distribution shown as violin plots and average values per embryo as dots. (**E, G**) For all graphs, E-cadherin intensity was normalized to the *WT* Center BEC group (Violin plots tested for two-tailed unpaired t test (Center Interlayer) and two-tailed Mann-Whitney test for all other comparisons, p<0.0001, n = 180 - 200 junctions (SEC and BEC junctions) and 90-100 cells (interlayer), dots n = 9 (*WT*) and 10 (*itgb1a*^+/-^*, itgb1b^+/-^*) embryos; 2 independent experiments). (**I, K**) For all graphs, desmoplakin intensity was normalized to the *WT* Center BEC group (Violin plots tested for two-tailed Mann-Whitney test, p<0.0001, n = 359 - 395 junctions (SEC and BEC junctions) and 188 - 190 cells (interlayer), dots n = 20 (*WT*) and 19 (*itgb1a*^+/-^*, itgb1b^+/-^*) embryos; 2 independent experiments).

Despite intact ECM composition, there was a profound decrease (p<0.0001) of the AJ component E-cadherin in both the center and periphery regions of the fin fold of *itgb1b^+/-^;itgb1a^+/-^*, not only between BECs but also at the BEC-SEC interface, and even between SECs (Fig. 3D-G). In contrast, desmoplakin levels increased in both center and periphery regions between BECs, SECs, and at the BEC-SEC interface in mutants compared to *WT* (Fig. 3H-K). Thus, mutants with reduced integrin β1 exhibit increased desmoplakin levels and decreased E-cadherin across the bilayer epidermis in both central and peripheral regions.

Given that SECs do not directly contact the ECM, these results suggest that reduced BEC–ECM adhesion has non-cell-autonomous effects on junctional organization in overlying SECs.

Together, these findings support a model in which integrin-mediated BEC–ECM interactions coordinate the balance of adherens and desmosomal junctions across both layers of the epidermis.

### ECM composition drives junctional specialization in a human epidermal model

Our data suggest that BEC-ECM interactions are required to differentially regulate cell-cell junctions throughout the bilayer epidermis. We hypothesized that the interactions between specific ECM proteins and BECs are sufficient to establish interlayer junctional specialization. To test this hypothesis, we employed a 3D *in vitro* cell culture model of bilayered human epidermis, enabling precise regulation of the BEC-ECM composition. Specifically, we engineered scaffold materials to mimic the central vs peripheral ECM in the fin fold.

Human keratinocytes cultured in low calcium conditions form a proliferative monolayer. When calcium levels are elevated, these keratinocytes undergo stratification within 24 hours (hrs), accompanied by the accumulation of terminal differentiation markers^42–44^. As expected, we found that treating human basal keratinocytes with Ca^2+^ for 48 hrs, resulted in two layers: basal keratinocytes positive for P63, and a layer of large suprabasal keratinocytes with significantly lower P63 expression, suggesting successful differentiation^45^ (Fig. 4A, B).

**Figure 4.**
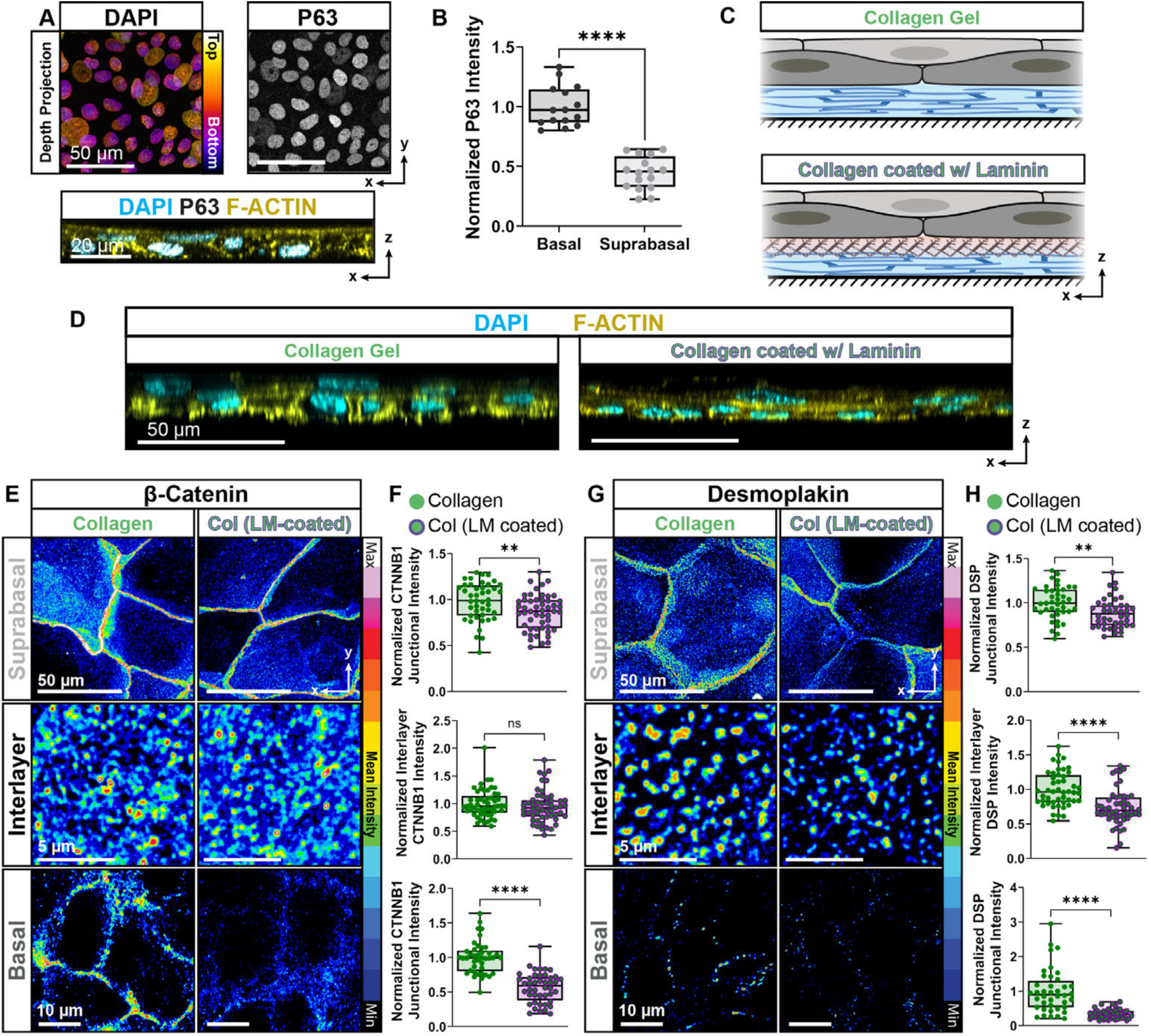
Human epidermal keratinocytes on different ECMs differentially regulate the expression of adherens junctions and desmosomes across layers *in vitro*. (**A**) Representative immunofluorescence images of P63, F-actin, and DAPI in both layers of the bilayer normal human epidermal keratinocytes (NHEK) cell culture system. DAPI image is z-depth color-coded. (**B**) Quantification of P63 intensity in Basal and Suprabasal cells (two-tailed unpaired t-test, p<0.0001, n=15 fields of view (FOV) containing 5-20 cells per FOV; 1 independent experiment). (**C**) Schematic of the two different gel constructs containing 100% collagen (Collagen) and 100% collagen gel coated with laminin (Col (LM-coated)). (**D**) Representative immunofluorescence z-stack cross sections of NHEKs cultured on each of the two gel systems. Cells were stained with F-actin and DAPI. (**E**) Representative immunofluorescence images of β-Catenin (CTNNB1) in culture on each gel type. Images are shown of the Suprabasal, Interlayer (between the Suprabasal and Basal), and the Basal cells. Immunofluorescence intensity is displayed using a 16-color scale. (**F**) Quantification of CTNNB1 intensity at Suprabasal junctions, Interlayer (between the Suprabasal and Basal), and Basal junctions in NHEKs cultured on the two gel systems. For each graph, intensity was normalized to the Collagen Gel Group (two-tailed unpaired t test (Suprabasal and Basal) and two-tailed Mann-Whitney test (Interlayer), p = 0.0046 (Suprabasal), 0.2213 (Interlayer), and p<0.0001 (Basal), n = 41-53 cells; 1 independent experiment). (**G**) Representative immunofluorescence images of Desmoplakin (DSP) in culture on each gel type. Images are shown of the Suprabasal, Interlayer (between the Suprabasal and Basal), and the Basal cells. Immunofluorescence intensity is represented as 16 colors. (**H**) Quantification of Desmoplakin intensity at Suprabasal junctions, Interlayer (between the Suprabasal and Basal), and Basal junctions in NHEKs cultured on the two gel systems. For each graph, intensity was normalized to the Collagen Gel Group (two-tailed unpaired t test (Suprabasal and Interlayer) and two-tailed Mann-Whitney test (Basal), p = 0.0042 (Suprabasal) and p<0.0001 (Interlayer and Basal), n = 40-50 cells; 1 independent experiment).

We seeded basal keratinocytes on 100% Collagen type I scaffold materials, the major component of the central fin fold and of the dermal ECM in humans, and on a Collagen type I scaffold coated with basement membrane Laminins (containing Laminin A, B1, and B2 chains) mimicking the peripheral fin fold (Fig. 4C). In both conditions, the addition of Ca^2+^ for 48 hrs still generated a bilyaered epidermis (Fig. 4D). Furthermore, we confirmed that basal keratinocytes produced the same types and levels of ECM under both conditions (Extended Figure 4A-D).

Despite these similarities, the junctional composition differed markedly by ECM scaffolds. Compared to basal keratinocytes seeded on Collagen-1 alone, those seeded on Laminin-coated collagen scaffolds displayed a profound decrease in the junction levels of the AJ component β-Catenin^46^ and of desmoplakin (Fig. 4E-H, Bottom panels). Suprabasal keratinocytes showed a similar, though less pronounced, Laminin-dependent reduction in β-Catenin and desmoplakin at junctions (Fig. 4E-H, Top panels). Notably, the Laminin-coated scaffold led to reduced levels of desmoplakin without affecting β-Catenin at the junctions between basal and suprabasal keratinocytes (Fig. 4E-H, Middle panels; Extended Data Fig. 4E-F).

These results suggest that ECM composition is sufficient to influence junctional specialization in bilayered epithelia. In particular, Laminins appear to selectively limit desmoplakin enrichment at interlayer contacts, mirroring patterns observed in the zebrafish fin fold.

### Laminin and collagen distinctly shape junctional architecture in the zebrafish epidermis

Our data thus far suggest that the differential distribution of ECM proteins in the fin fold creates distinct intercellular junctions in the bilayer epidermis. To assess the physiological role of specific ECM components, we generated zebrafish mutants that disrupt laminin a5 (*lama5*) and collagen type I alpha 1 chain (*col1a1a*), two of the most enriched ECM proteins in the BEC (Extended Data Fig. 1F).

Homozygous mutants for both *lama5* and *col1a1a* display abnormal fin development ^25,47^, so we used heterozygous embryos, which maintain normal development and intact fin folds, enabling us to examine bilayer architecture (Extended Data Fig. 5A). Heterozygous mutants exhibited reduced levels of their respective ECM proteins throughout the fin fold, with *lama5^+/-^* showing a prominent reduction of laminin a5 at the periphery and *col1a1a*^+/-^ showing a reduction of collagens in the central zone (Fig. 5A-C).

**Figure 5.**
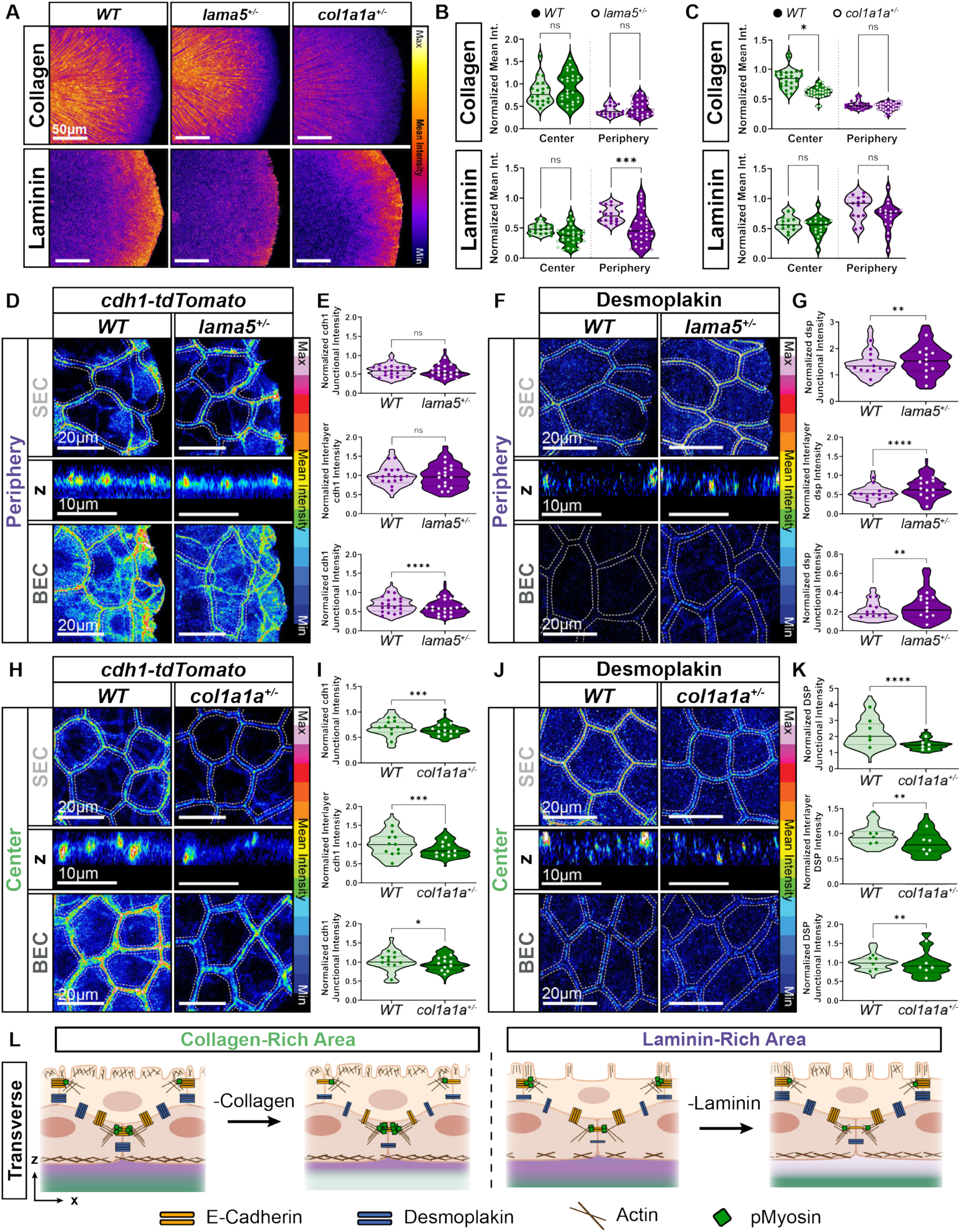
BEC-Laminin Interactions Regulate Desmosomes, AJs, and Actomyosin Signaling in the Peripheral Bilayer Epidermis of the Fin Fold. (**A**) Representative maximum projection of *wild type* (*WT*) vs. *lama5^+/-^ vs. col1a1a^+/-^* fin fold, stained with a FAM-tagged collagen hybridizing peptide and Laminin A1 at ∼48 hpf. Intensity displayed using fire LUT. (**B)** Quantification of average normalized collagen and laminin intensity per embryo. (Collagen: Kruskal-Wallis test with multiple comparisons, p>0.9999, n = 20 (*WT*) and 26 (*lama5^+/-^*) embryos; 2 independent experiments; Laminin: Ordinary One-Way ANOVA test with multiple comparisons, p = 0.2249 (Center) and 0.0006 (Periphery), n = 18 (*WT*) and 28 (*lama5^+/-^*) embryos; 2 independent experiments. (**C**) Quantification of average normalized collagen and laminin intensity per embryo. (Collagen: Kruskal-Wallis test with multiple comparisons, p = 0.0246 (Center) and >0.9999 (Periphery), n = 27 (*WT*) and 33 (*col1a1a ^+/-^*) embryos; 3 independent experiments; Laminin. Ordinary One-Way ANOVA test with multiple comparisons, p = 0.8872 (Center) and 0.2019 (Periphery), n = 13 (*WT*) and 18 (*col1a1a ^+/-^*) embryos 2 independent experiments. (**D, F**) Representative (**D**) images of *cdh1-tdTomato^xt^*^18^ and (**F**) immunofluorescence staining for desmoplakin in the peripheral region of zebrafish fin folds at ∼48 hpf.. Z-stack images taken from the SEC layer to the BEC layer. SECs and BECs were identified by maximum projection of selected z slices based on their z-position. Representative interlayer images are shown in orthogonal view of single z slice. Intensity displayed using 16-color scale. (**E, G**) Quantification of (**E**) E-cadherin and (**G**) desmoplakin at peripheral SEC junctions, interlayer (between SEC and BEC), and BEC junctions, with individual junctional measurements distribution shown as violin plots and average values per embryo as dots. (**E**) For all graphs, E-cadherin intensity was normalized to the *WT* Center BEC group in panel Extended Data Fig. 5C (two-tailed Mann-Whitney test on violin plots, p = 0.2132 (SEC), 0.4736 (Interlayer), and p<0.0001 (BEC), n = 348 - 363 junctions (SEC and BEC junctions) and 160-169 cells (interlayer), dots n = 19 (*WT*) and 18 (*lama5^+/-^*) embryos; 3 independent experiments). (**G**) For all graphs, desmoplakin intensity was normalized to the *WT* Center BEC group in panel Extended Data Fig. 5E (two-tailed Mann-Whitney test on violin plots, p = 0.0014 (SEC), 0.0036 (BEC), and p<0.0001 (Interlayer), n = 232 - 240 junctions (SEC and BEC junctions) and 117-120 cells (interlayer), dots n = 12 (*WT*) and 12 (*lama5^+/-^*) embryos; 2 independent experiments) (**H, J**) Representative (**H**) images of *cdh1-tdTomato^xt^*^18^ and (**J**) immunofluorescence staining for desmoplakin in the central region of zebrafish fin folds at ∼48 hpf. Images show SECs, Interlayer (between SEC and BEC), and BECs. Confocal z-stack images were taken from the SEC layer to the BEC layer. SECs and BECs were identified by maximum projection of selected z slices based on their z-position. Representative interlayer images are shown in orthogonal view of single z slice. Intensity displayed using 16-color scale. (**I, K**) Quantification of (**I**) E-cadherin and (**K**) Desmoplakin at central SEC junctions, interlayer (between SEC and BEC), and BEC junctions, individual junctional measurement distribution shown as violin plots and average values per embryo as dots. (**I**) For all graphs, E-cadherin intensity was normalized to the *WT* Center BEC group in bottom panel Fig. 5I (two-tailed unpaired t test for BEC Center and two-tailed Mann-Whitney test on all other violin plots, p = 0.0001 (SEC), 0.0009 (Interlayer), and 0.04825 (BEC), n = 170 junctions (SEC and BEC junctions) and 110 cells (interlayer), dots n = 11 (*WT*) and 11 (*col1a1a^+/-^*) embryos; 1 independent experiment). (**K**) For all graphs, Desmoplakin intensity was normalized to the *WT* Center BEC group in panel Fig. 5K (two-tailed unpaired t test on Interlayer, and two-tailed Mann-Whitney test on all other violin plots, p< 0.0001 (SEC), p = 0.01437 (BEC) and 00011 (Interlayer); n = 62-82 junctions (SEC and BEC junctions) and 50-60 cells (interlayer), dots n = 6 (*WT*) and 6 (*col1a1a^+/-^*) embryos; 1 independent experiment) (**L**) Summary schematic of junctional characteristics in the peripheral regions of SECs, SEC-BEC interface, and BECs.

First, we analyzed *lama5^+/-^* mutant fin folds. For both the central and peripheral regions, E-cadherin levels were lower between BECs, but not between SECs or the SEC-BEC interface, in *lama5+/-* compared to WT (Fig. 5D and E). By contrast, desmoplakin levels were elevated throughout the *lama5*^+/-^ mutant versus WT: between BECs, SECs, and BEC-SECs, in the central and peripheral regions (Fig. 5F and G) (Extended Data Fig. 5B-I).

Next, we examined *col1a1a⁺/⁻* mutants. In the central region, we observed a significant reduction in the levels of E-cadherin and desmoplakin across all layers in the mutant compared to *WT* (Fig. 5H-K). E-cadherin and desmoplakin junction levels were both decreased in the peripheral region of *col1a1a⁺/⁻* fins in BECs, BEC-SECs but not in SECs compared to WT (Extended Fig. 5J-Q).

Together, these findings suggest that BEC-laminin interactions promote AJ formation while limiting desmosomal enrichment, whereas BEC–collagen interactions support both junction types across layers, particularly in the central fin fold (Fig. 5L).

### Region-specific ECM requirements for wound closure in the zebrafish epidermis

SECs within the periderm in both humans and zebrafish play a critical role in protecting embryonic BECs against external physical forces. We hypothesized that BEC-laminin interactions, and in turn the differential adhesion profiles in the central versus peripheral periderm, might affect its response to mechanical injury. To test this hypothesis, we used a two-photon laser to induce a single-cell ablation in the SEC layer.

Strikingly, SECs in the laminin-enriched peripheral region completed wound closure faster than SECs in the collagen-enriched center region (Fig. 6A and B, Supplementary Video 2). These observations establish a region-specific difference in epithelial repair and suggest that the ECM composition may influence the wound healing response.

**Figure 6.**
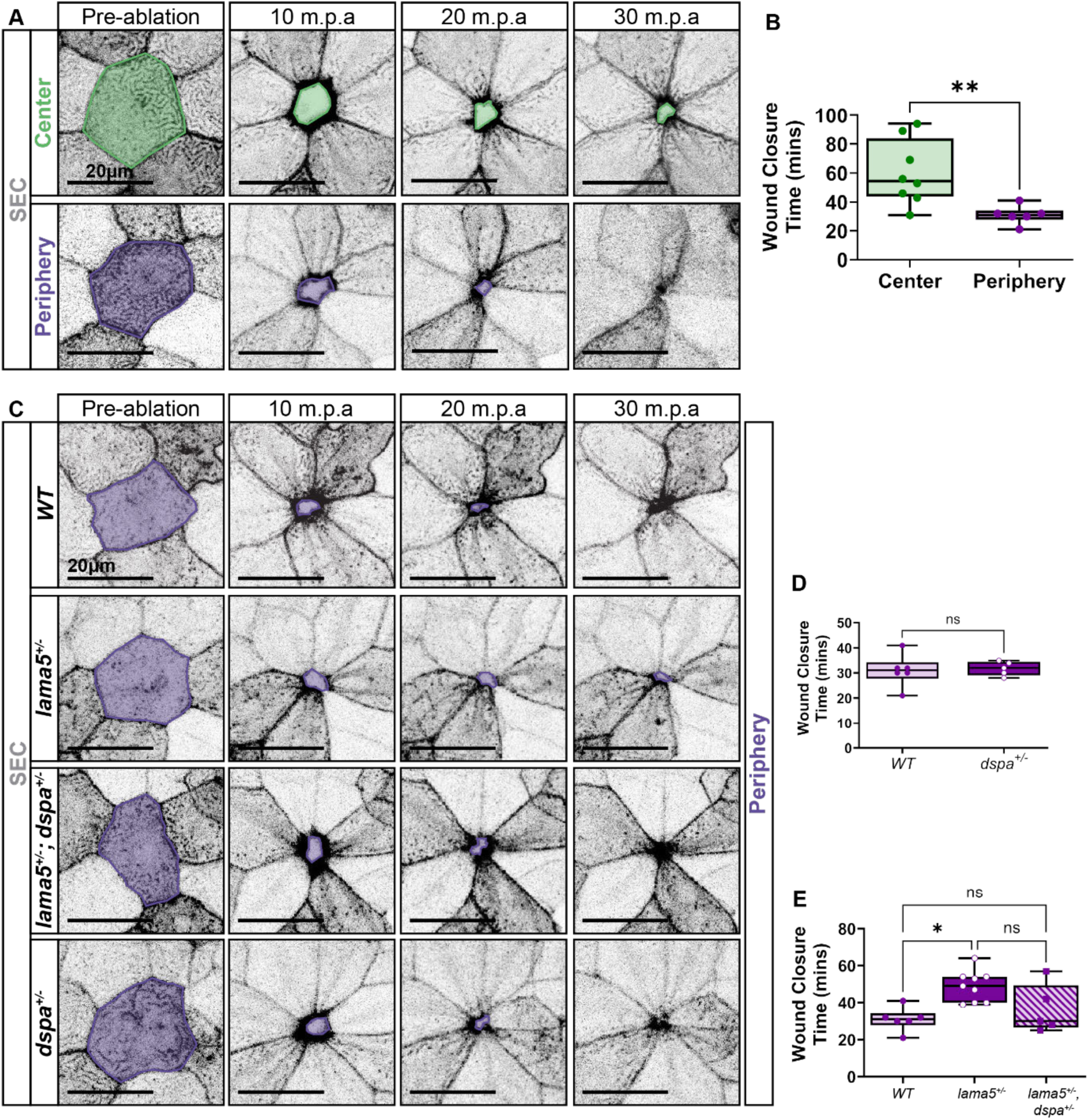
BEC-Laminin Interactions are Required for Efficient Wound Closure in the Periderm. (**A)** Representative live imaging confocal images of *Tg(krt5:Gal4ff)^la2^*^12^*; Tg(UAS:LifeAct-GFP)^mu2^*^71^ for LifeAct-GFP under the SEC-specific *krt5* promoter, showing the central and peripheral regions of the fin fold at 2 dpf before and after single-cell ablation. Shaded border indicates the ablated cell and subsequent wound closure. (**B**) Quantification of time (min) required for complete wound closure. Box plots with min to max, each point representing indicated measurement for each ablation experiment. (two-tailed Mann-Whitney test, p = 0.0040, n = 8 (Center) and 6 (Periphery) embryos; 4 independent experiments). (**C)** Representative live imaging confocal images of *Tg(krt5:Gal4ff)^la2^*^12^*; Tg(UAS:LifeAct-GFP)^mu2^*^71^ for LifeAct-GFP under the SEC-specific *krt5* promoter in the peripheral regions of fin fold of indicated genotypes at 2 dpf before and after single-cell ablation. Shaded border indicates the ablated cell and subsequent wound closure. (**D)** Quantification of periphery wound closure time (min), comparing *WT* and *dspa^+/-^* mutants. Box plots with min to max, each point representing indicated measurement for each ablation experiment. (Mann-Whitney test, p = 0.7879, n = 6 (*WT*) and 5 (*dspa^+/-^*) embryos; 4 independent experiments) (**E**) Quantification of wound closure time (min), comparing periphery *WT*, *lama5^+/-^*, and *lama5^+/-^, dspa^+/-^* mutants. Box plots with min to max, each point representing indicated measurement for each ablation experiment (Kruskal-Wallis test with multiple comparisons; p = 0.0273 (*WT* vs. *lama5^+/-^*), >0.9999 (*WT* vs. *lama5^+/-^, dspa^+/-^*), and 0.1988 (*lama5^+/-^* vs. *lama5^+/-^, dspa^+/-^*); n = 6 (*WT*), 9 (*lama5 ^+/-^*), and 5 (*lama5^+/-^, dspa^+/-^*) embryos; 5 independent experiments.

To examine the role of laminin more directly, we assessed wound healing in *lama5^+/−^* mutants. SECs took longer to complete wound closure after single-cell injury in the peripheral region of the *lama5^+/−^* mutant fin fold compared to *WT*, whereas repair time in the center region was unaffected (Fig. 6C, E, and Extended Data Fig 6A, B; Supplementary Video 3, 4).

Moreover, compared to *WT*, *lama5^+/−^* peripheral SECs had increased levels of microridge actin, peripheral BECs had elevated cortical actin organization, and both peripheral cell layers had reduced junction levels of phosphorylated myosin (Extended Fig. 6C-H). These changes were not observed in the central region of *lama5^+/−^* versus WT (Extended Fig. 6I-N). These findings indicate that laminin a5 is required in the periphery to support efficient wound closure, likely through regulation of actin localization and contractility.

Next, we assessed wound closure in *col1a1a^+/−^* fins. In contrast to the *lama5^+/−^* mutant fin fold, the time required for peripheral SECs to complete wound healing was comparable in *col1a1a^+/−^*and *WT* (Extended Data Fig 7A, B, Supplementary Video 5), whereas centrally positioned SECs took longer to close wounds in *col1a1a^+/−^* fin folds compared to *WT* (Extended Data Fig 7C, D, Supplementary Video 6). These findings suggest that collagen plays a more prominent role in promoting repair within the center of the fin fold epithelium.

Because BEC-laminin interactions limit, whereas BEC-collagen interactions promote, desmoplakin expression, we hypothesized that the increased desmoplakin in the periphery of *lama5^+/−^* fin folds underlies impaired wound closure. To test this, we reduced desmoplakin levels by generating *dspa* mutants. Zebrafish desmoplakin is encoded by two paralogs, *dspa and dspb,* with *dspa* being the predominant isoform expressed in BECs (Extended Data Fig. 7E-G). We analyzed the fin fold in heterozygotes to avoid the gross morphological defects reported for homozygous mutants^48^, and observed a small but significant reduction of desmoplakin protein levels in peripheral but not central BECs and SECs of *dspa^+/-^* mutants (Extended Data Fig. 7H-M). We found that the time to complete wound closure was similar in *dspa^+/−^*mutant and *WT* peripheral fin folds (Fig. 6C, D and Supplementary Video 7). Importantly, *lama5^+/−^; dspa^+/−^*double mutants showed partially rescued wound closure times compared to *lama5^+/−^*mutants, comparable to *WT* (Fig. 6 C,E, Supplementary Video 8). These results suggest that elevated desmoplakin and desmosome accumulation in a laminin-deficient background restricts wound repair to the periphery region.

Overall, our findings suggest that regional ECM composition tunes the wound-healing capacity of the zebrafish fin fold epidermis through its influence on BECs and, in turn, junctional architecture and cytoskeletal organization (Fig. 7).

**Figure 7.**
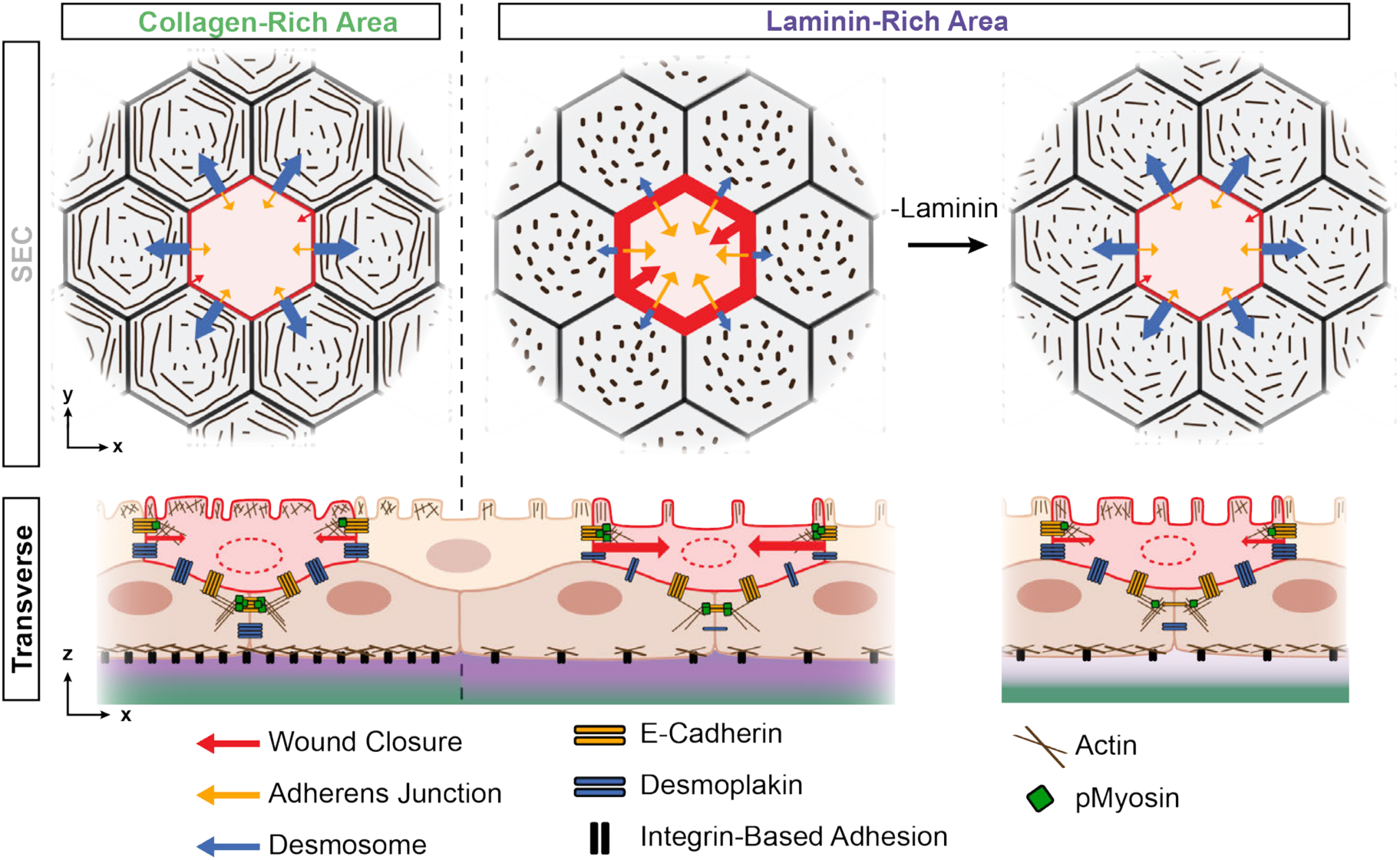
Schematic Illustration of how BEC-ECM Interactions Influence Periderm Architecture and Wound Repair. We propose that the interaction between basal epidermal stem cells (BECs) and laminin influences the junctional organization and injury response of the periderm, via the regulation of interlayer junction dynamics. The developing bilayered skin appendage is organized as collagen-enriched areas, where BECs exhibit increased adherens junction (AJ) actomyosin signaling and desmosome formation. Superficial epidermal cells (SECs) layer (the periderm) above these BECs display reduced motility, characterized by stronger desmosomal junctions and less actomyosin signaling. In laminin-rich areas, BECs show decreased actomyosin activity and reduced AJ and desmosome formation within the intralayer, but they unexpectedly maintain interlayer AJ and repress desmosome formation. SECs above these BECs have faster motility during wound closure. Partial removal of laminin matrices further reduces BEC actomyosin activity, AJ and desmosome formation while increasing desmosome connections at the interfaced membrane with SECs. SECs, connected to these BECs, exhibit increased desmosome overall, and slow wound closure.

## Discussion

Here, we identify a region-specific adhesion profile within the periderm in the fin fold acquired through interactions between basal epidermal stem cells (BECs) and the extracellular matrix (ECM). The ECM is known to regulate BECs differentiation and fate^23,49–54^. Our findings provide, to our knowledge, the first evidence that BEC-ECM interactions influence injury resilience of an interconnected tissue without affecting their differentiation into stratified cell layers. During embryonic development, superficial epidermal cells (SECs) form a protective layer, the periderm, that extends from the developing limb (fin) to other regions of the embryo body and plays a crucial role in shielding the underlying stem cells from mechanical stresses. We found that the response of the periderm of the fin fold to cell injury is directly dependent on BEC-ECM interactions. BECs organized their junctions with SECs in a region-specific manner, forming both desmosomes and AJs above the central, collagen-enriched ECM, but predominantly AJs above the peripheral, laminin-enriched ECM. We propose that BEC-laminin interactions serve to suppress desmosomes across layers, this shift decreases cellular resistance while preserving SEC actomyosin contractility and tissue integrity, enabling the SECs above the BEC-laminin areas to close wounds faster following cell injury (Fig. 7).

The regulation of cell-cell junctions and cell-ECM interactions has been extensively studied in the context of cell injury and wound healing^20,55,56^. A profound reorganization of actomyosin contraction typically occurs during wound closure^57^, and additional junctional forces, including cell resilience generated by desmosomes at the junction can be activated due to substrate stiffness or cell deformation^58–62^. Overall junctional tension is enhanced through desmosomes and adherens junctions during tissue injury responses. Consequently, the absence of desmosomes is often linked to impaired wound healing, loss of tissue integrity and the formation of blistering in epidermis^63^. In contrast, our data suggest that interactions between BECs and laminin-rich matrices suppress interlayer desmosomes and while maintaining the contacts with SEC via AJ, desmosomes repression promote wound closure. This apparent inconsistency may stem from several factors. First, most cell ablation studies have focused on monolayer cell models in direct contact with the ECM, where actin cytoskeleton reorganization must coordinate junctional and focal adhesion tensions to support both actomyosin contraction and cell migration. In contrast, cell injury within stacked cell layers—such as the bilayered epidermis—has, to our knowledge, never been directly evaluated in the context of inter-tissue interactions. Second, based on our model, we propose that desmosomes at the SEC-BEC interface enhance the resistance to cell deformation^64^ triggered by the death of SECs. Hence, fewer desmosomes at the SEC-BEC interlayer reduce SEC resistance and accelerate wound repair. This is in agreement with the dynamic needs of the developing fin fold, where peripheral regions also have less desmosome presence. This arrangement may balance the simultaneous expansion of skin stem cells and their protection by the periderm during skin repair at the appendages. Despite this reduction in desmosomes, we do not observe increased interlayer movement during wound closure or development, likely because AJs and cortical actin networks maintain coordinated coupling between BECs and SECs. Thus, reduced desmosome levels may enhance cellular dynamics without compromising overall interlayer integrity.

The organization of the ECM plays a crucial role in determining the distribution and type of junctions in monolayered cells, mediated by interactions with various cytoskeletal proteins. ECM distribution is also generally critical for organ morphogenesis. For instance, laminin typically supports basement membrane stability and epithelial integrity, but it is also required to promote osteogenesis^25,65,66^. Hence, full removal of laminin a5 abolishes fin development^25^ and thus its effect on epithelial integrity can be tested only with partial loss of function models.

Similarly, collagen-rich matrices provide stiffness and tensile strength and are key for mesenchymal growth during the initial limb development^26,47,67^. Collagen is produced by both basal stem cells and embryonic fibroblasts, which are enriched in the collagen-dense central areas of the fin fold. In contrast, laminins are primarily produced by basal epidermal stem cells^27^ and are concentrated at the fin’s very tip, where fibroblasts are sparse. Since complete deletion of these ECM proteins severely disrupts fin growth, and tissue-specific deletion remains to be investigated, we acknowledge that our data only correlate BEC–ECM interactions with effects on SECs. Thus, other cell types may also be necessary to confer laminin-BEC specialization within the fin fold. Nevertheless, our *in vivo* studies suggest that central BECs on collagen-enriched areas exhibit increased AJ actomyosin signaling and desmosome formation compared to BECs on laminin-enriched areas. Notably, peripheral BECs in *lama5*^+/-^ mutants, which interact with reduced levels of laminin, exhibit a different adhesion molecule organization compared to *WT* central BECs, which also interact with relatively low laminin levels. This suggests that collagen alone in the peripheral regions with decreased laminin is insufficient to recapitulate the BEC architecture observed in the *WT* center. These differences could also be attributed to ECM properties beyond protein composition, such as the architecture and stiffness of the basement membrane, or other microenvironmental differences such as tissue signaling functions. Further investigation is needed to understand how these properties differentially influence surrounding tissues through their interactions with stem cell compartments.

Communication between skin layers has been observed in both zebrafish embryo (bilayered skin) and multilayered mouse skin^10,22,37^. For instance, the loss of E-cadherin function in either the periderm or BECs affects E-cadherin levels in the other layer. E-cadherin functions as a transducer of polarity between the periderm and BECs^22^. It regulates the localization and levels of Lgl2, a basolateral polarity regulator, in both a layer-autonomous and non-autonomous manner. Lgl2 plays a critical role in the formation of hemidesmosomes, which mediate basal cell-to-matrix adhesion through integrin α6β4^37^. Since hemidesmosomes bind primarily to laminin, we currently do not know whether hemidesmosomes play a role in maintaining E-cadherin basolateral polarity in the periphery. However, hemidesmosomes have not formed at 2 dpf^37^. In mice, intratissue tension generated by the upper epidermal layers through microtubule regulation significantly impacts the behavior of BECs. Increased contractility in these differentiated cells, which can result from microtubule disruption, promotes stem cell hyperproliferation while inhibiting their migration. This highlights that stem cells in stratified tissues respond not only to substrate rigidity but also to mechanical cues from intratissue tension, with differentiated cells playing a critical role in shaping stem cell behavior and maintaining tissue integrity^10^.

Overall, our findings reinforce existing evidence of intratissue communication and propose a new paradigm for embryonic epidermal stem cells. We propose that periderm near limb appendages have enhanced wound healing properties based on regionally-specific differences in the underlying stem cell-matrix interactions. Possibly by secreting distinct ECMs, and regulating adhesion molecule expression, epidermal stem cells regulate the protective function of the interconnected, differentiated tissue, the periderm. This discovery holds significant implications for our understanding of BEC contributions during epidermal development, and can inform future applications such as tissue engineering and regenerative medicines, including organ repair during and skin transplants.

## Methods

### Zebrafish Husbandry

Zebrafish were raised and maintained at 28.5°C using standard methods and according to protocol approved by the Yale University Institutional Animal Care and Use Committee (#2020-11473). AB strain zebrafish (ZDB-GENO-960809-7) were used as wild-type genetic background. The following zebrafish transgenic lines have been previously described: *TgBAC(ΔNP63:Gal4ff)^la2^*^13^ (ZDB-ALT-150424-4)^68^, *Tg(krt5:Gal4ff)^la2^*^12^ (ZDB-ALT-150424-3)^68^, *Tg(UAS:NLS-GFP)^el6^*^09^ (ZDB-ALT-210709-6)^69^, *Tg(krt5:LifeAct-Ruby)^la2^*^27^ (ZDB-ALT-200226-18)^38^, *Tg(UAS:LifeAct-GFP)^mu2^*^71^ (ZDB-ALT-130624-2)^70^, *TgBAC(pdgfrb:eGFP)^ncv22Tg^* (ZDB-ALT-160609-1)^71^, *Tg(Bactin:HRAS-EGFP)^vu1^*^19^ (ZDB-ALT-061107-2)^72^, and *TgBAC(lamC1:lamC1-sfGFP)^sk1^*^16^ (ZDB-TGCONSTRCT-241010-2)^31^. The following endogenous knock-in lines have been previously described: *cdh1-tdTomato^xt^*^18^ (ZDB-ALT-190419-2)^73^ and *itgb1b:itgb1b-sfGFP^sk1^*^33^ (ZDB-ALT-241011-7)^31^. The zebrafish mutants used in this study have been previously described: *itgb1a^sk1^*^09^ (ZDB-ALT-240116-1) and *itgb1b^sk1^*^10^ (ZDB-ALT-240116-2)^31^. The *lama5^ya3^*^77^, *col1a1a^ya3^*^92^, and *dspa^ya3^*^91^ mutant lines were generated in this study.

### Generation and Genotyping of Mutant Alleles

To generate *lama5^ya^*^37^, *col1a1a^ya3^*^92^, and *dspa^ya3^*^91^ mutants, we followed previously described CRISPR-Cas9-based gene-editing protocols^74^. CRISPRScan (https://www.crisprscan.org/) was used to design guide RNAs (gRNAs) to generate loss-of-function mutant of *lama5.* gRNA preparation was performed as previously described^75^. Briefly, WT embryos were injected with 100 pg of gRNAs and 200 pg of *Cas9* mRNA at the one-cell stage. PCR genome amplification and T7E1 assay was used to validate indels as previously^75^.

The *lama5^ya3^*^77^ mutant contains 23-bp deletion in *lama5* exon2 that leads to a pre-mature stop codon in exon3. *lama5^ya3^*^77^ homozygous mutant embryos display complete loss of the fin fold, which is consistent with previous reports^25^. Sequences and primers are listed in supplementary Table 1. To genotype *lama5^ya3^*^77^, zebrafish embryos and adults were genotyped using GeneMarker (v2.4.0) as previously described^76^, with genotyping primers listed in supplementary Table 1.

The *col1a1a^ya3^*^92^ mutant contains 5-bp deletion in *col1a1a* exon2 that leads to a pre-mature stop codon in exon3. *col1a1a^ya3^*^77^ homozygous mutant embryos display damaged fin fold, which is consistent with previous reports^47^. Sequences and primers are listed in supplementary Table 1. To genotype *col1a1a^ya3^*^92^, zebrafish embryos and adults were genotyped using GeneMarker (v2.4.0) as previously described^76^, with genotyping primers listed in supplementary Table 1.

The *dspa^ya3^*^91^ mutant contains 5-bp deletion in *dspa* exon2 that leads to a pre-mature stop codon in exon3. Sequences and primers are listed in supplementary Table 1. To genotype *dspa^ya3^*^91^, zebrafish embryos and adults were genotyped using GeneMarker (v2.4.0) as previously described^76^, with genotyping primers listed in supplementary Table 1.

To genotype *itgb1a^sk1^*^09^ and *itgb1b^sk1^*^10^, zebrafish embryos and adults were genotyped by PCR as previously described^31^, with genotyping primers listed in supplementary Table 1.

### Zebrafish Embryos Immunofluorescence Staining

#### Immunofluorescence

Immunofluorescence (IF) was performed as follows for all zebrafish embryos. After overnight 4% paraformaldehyde (Santa Cruz) fixation at 4°C, 48 hpf embryos were washed with 1X PBS-0.01%Tween-20 (PBSTw) four to five times for 5 minutes, and then permeabilized with 0.125% trypsin (Millipore Sigma T4549) for 5 minutes. Embryos were washed in blocking solution (0.8% Triton-X, 10% normal goat serum, 1% BSA, 0.01% sodium azide in PBSTw) four times for 5 minutes, followed by an additional two-hour incubation in blocking solution with shaking at room temperature. Primary and secondary antibodies incubation were conducted overnight at 4°C. Following each overnight antibody incubation, six washes for a total of 4 hours were performed with washing buffer (1% Triton-X in PBSTw) at room temperature. Primary antibodies used were specific to RFP (1:250, Antibodies Online, ABIN129578), Laminin α1 (1:250, Sigma-Aldrich, L9393), phosphorylated Myosin Light Chain (Ser19) (1:200, Cell Signaling, Cat. #3675), Desmoplakin (undiluted, ProGen, 651155), and Vinculin (1:100, Sigma-Aldrich, V9131). Secondary antibodies for either rabbit (1:250, ThermoFisher, A10040) or mouse (1:250, ThermoFisher, A21202) were used. To stain nuclei, DAPI (1:500, ThermoFisher, D1306) was added during the secondary antibody incubation step. To stain F-actin, Phalloidin-568 (1:100, Invitrogen, A12380) was added during the primary antibody incubation step. Samples were mounted in 1% Low Range Ultra Agarose (Bio-Rad, #1613106) and imaged using the Zeiss LSM980 upright confocal microscope (ZEN 3.4, blue edition, Carl Zeiss) equipped with a water-immersed x10/0,3 or 20/0,5 or x40/1,0 objective lens.

#### Collagen Hybridizing Peptide

To stain denatured collagen, Collagen hybridizing peptide with 5-FAM or Cy3 conjugate (5 µM, F-CHP/ R-CHP, 3Helix, FLU60/ RED60) was used. After overnight 4% paraformaldehyde (Santa Cruz) fixation at 4°C, 48 hpf embryos were washed with 1X PBS-0.01%Tween-20 (PBSTw) four to five times for 5 minutes, and then permeabilized with 0.125% trypsin (Millipore Sigma T4549) for 5 minutes. Embryos were washed in blocking solution (0.8% Triton-X, 10% normal goat serum, 1% BSA, 0.01% sodium azide in PBSTw) four times for 5 minutes, followed by an additional two-hour incubation in blocking solution with shaking at room temperature. The CHP stock solution (50 µM) was diluted in PBS buffer to make the working solution (5 µM) and heat-activate by incubating at 80°C for 5 minutes. The solution was immediately cooled on ice for 30-90s, to quench it to room temperature, then quickly applied to fixed embryos. Laminin α1 (1:250, Sigma-Aldrich, L9393) primary antibody was added to the diluted CHP solution before applying to the samples. Samples were incubated with the staining solution at 4°C overnight with shaking. Following each overnight antibody incubation, six washes for a total of 4 hours were performed with washing buffer (1% Triton-X in 1x PBSTw) at room temperature.

### Cell Immunostaining

For immunofluorescence analysis in NHEKs, cells were fixed with 4% paraformaldehyde (Electron Microscopy Sciences) for 20 minutes. Cells were washed with DPBS with calcium and magnesium (Thermo, 21300025) and permeabilized in 0.5% Triton X-100 for 10 minutes. Cells were blocked with 10% serum for 1 hour, incubated for either 1 hour at room temperature or overnight at 4°C with primary antibodies, and incubated for 1 hour with secondary antibodies.

Primary antibodies used were specific to p63 (1:100, BioCare Medical, CM163A), β-catenin (1:500, Sigma, C2206), Desmoplakin (undiluted, ProGen, 651155), Human Collagen Type I (1:50, CedarLane, CL50111AP-1), Collagen Type IV (1:100, ThermoFisher, 19674-1-AP), and Laminin (1:500, ThermoFisher, PA5-143904). Secondary antibodies for either rabbit (1:1000, ThermoFisher, A-32731 or A-21244), mouse (1:1000, ThermoFisher, A-11001 or A-21235), or Rat (1:1000, ThermoFisher, A-11006 or A-21247) were applied. To stain F-actin, phalloidin (1:400, ThermoFisher, A12380) was added during the secondary antibody incubation step.

Samples were mounted with DAPI in Fluoromount-G (Thermo, 00-4959-52) and imaged on Zeiss LSM980 Airyscan 2 confocal.

### Confocal Live and Time Lapse Imaging of Zebrafish Embryos

Zebrafish embryos were treated with 0.003% 1-phenyl-2-thiourea (PTU, MilliPore Sigma, P7629) starting at 70%/80% gastrulation stage to prevent pigmentation. Embryos imaged live by confocal microscopy were anaesthetized in 0.1% tricaine and mounted in 1% (w/v) Low Range Ultra Agarose (Bio-Rad, #1613106) at 48hpf. Confocal live imaging was performed using the Zeiss LSM980 upright confocal microscope (ZEN 3.4, blue edition, Carl Zeiss) equipped with a water-immersed x10/0,3 or 20/0,5 or x40/1,0 objective lens. Z-stacks were acquired at 0.5 µm increments. Confocal time-lapse movies were recorded for wild type embryos in *Tg(krt5:Gal4ff)^la2^*^12^*, Tg(UAS:LifeAct-GFP)^mu2^*^71^*; cdh1-tdTomato^xt^*^18^ background, at 28°C incubation stating at 48 hpf. Z-stacks were acquired at 2 µm increments with intervals of every 10 min for a total duration of 8 hours. For each confocal imaging session, the laser power was consistently set to the same level across all samples. Cells were tracked using dragon-tail analysis in Imaris software (V.9.9.1, Bitplane).

### Two-Photon Single-Cell Laser Ablation and Time Lapse Imaging

Single cell laser ablation of superficial epidermal cells (SECs) was performed using a Leica Stellaris 8 DIVE multiphoton/confocal microscope equipped with a Spectra Physics MaiTai laser at 2 dpf. *Wild type* and *lama5^+/-^*embryos, in the *Tg(krt5:Gal4ff)^la2^*^12^*, Tg(UAS:LifeAct-GFP)^mu2^*^71^ background, were anaesthetized in 0.1% tricaine and mounted in 1% (w/v) Low Range Ultra Agarose (Bio-Rad, #1613106). Prior to ablation, confocal z-stacks were set up at 2 µm increments to capture pre-ablation images. Laser ablation was performed in a 10 µm diameter region of interest with the MaiTai laser at a wavelength of 810 nm with the laser power at 70% for two iterations of 0.743 seconds. Confocal z-stack images were acquired at 1-minute intervals until wound closure.

### Cell Culture

Normal human epidermal keratinocytes (NHEKs) were purchased from Promo Cell (C-12001) from a 4-year-old Caucasian male donor (Lot #: 494Z030.1). Cells were screened by manufacturer for bacteria, yeast, fungi, mycoplasma, virus and authenticated for positive expression of cytokeratin via flow cytometry. NHEKs were cultured at 37°C, 5% CO_2_, and 95% relative humidity according to manufacturer’s instructions up to passage four. Unless otherwise specified, cells were cultured in Keratinocyte Growth Medium 2 (PromoCell, C-20111) which contains no serum, low CaCl_2_ (0.06 mM), and supplemented with Gentamicin (Thermo, 15710064) at a final concentration of 5 μg/mL.

### Production of collagen and laminin gels

Collagen and laminin gels were formed on 35mm glass-bottom dishes (MatTek, P35G-0-20-C) by first coating dishes with 1.8 mg/mL polydopamine coating solution made from dopamine-HCL (Acros Organics, AC122000100) in 10mM Tris-HCL (pH 8.5). Dopamine solution was rinsed 3 times with ddH_2_O, the coated glass was fully dried in a warm oven overnight and cooled to room temperature the next day. To make the neutralized collagen solution (2 mg/mL final concentration of collagen), sterile ddH_2_O, 10x PBS (Corning, 46-013-CM), Collagen (Corning, 354249), and NaOH were mixed together. All solutions and tubes used for making gels were pre-chilled on ice. Next, 23 uL of the neutralized collage was dispensed onto the dopamine-HCL coated glass and an autoclaved 8 mm round coverslip (Electron Microscopy Sciences, 7229608) was placed on top of the droplet to ensure even coating. The collagen solution was allowed to solidify in an incubator at 37C for 40 minutes. After full gelation, the gels were soaked with PBS for 15 minutes and the top coverslip was removed. To make the laminin coated collagen gel, collagen gels were created as described above. A 50%, ∼5mg/mL laminin (Corning, 354259, 10 mg/mL) solution was made in ice-cold PBS and 20 uLs were evenly dispensed on top of the collagen cushion. The laminin layer was allowed to cure at 37C for 20 minutes. After curing, gels were inspected using a phase microscope to ensure the laminin layer adhered to the collagen gel. Gels were stored in PBS until seeding.

### NHEK stratification on collagen and laminin gels

NHEKs were expanded in T-75 flasks (ThermoFisher, 156499) and lifted into a single cell suspension using Trypsin-EDTA (0.05%) with phenol red (ThermoFisher, 25300054). To obtain confluent monolayers on gels within 24 hours of seeding, 800,000 cells were seeded into each dish. Cells were seeded and grown overnight in Keratinocyte Growth Medium 2 with low CaCl_2_ (0.06 mM). 24 hours after initial seeding, media was changed to Keratinocyte Growth Medium 2 supplemented with high CaCl_2_ (1.2mM). Cells were grown for 48 more hours with a media change every 24 hours. Cells were then fixed in 4% paraformaldehyde (Electron Microscopy Sciences) for 20 minutes before proceeding with immunofluorescence staining.

### Image Analysis and Quantifications

#### Zebrafish Confocal Fluorescence Images

Confocal fluorescence images were analyzed with Imaris software (V.9.9.1, Bitplane, Oxford Instruments) and the Fiji version of ImageJ (NIH)^77^. P63 expression across the fin fold was quantified using live confocal images of *TgBAC(ΔNP63:Gal4ff)^la2^*^13^*; Tg(UAS:NLS-GFP)^el6^*^09^. Each basal epidermal stem cell nucleus (GFP+) was 3D-reconstructed using Imaris surface module based on fluorescence intensity. Mean intensity and x,y,z coordinates (µm) for each BEC nucleus were exported for statistical tests. Distance from the center for each BEC nucleus was calculated using the x,y,z coordinates, and mean intensity was normalized to each embryo’s average intensity.

Confocal z-stack images were taken from the SEC layer to the BEC layer. SECs and BECs were identified by maximum projection of selected z slices based on their z-position. Intralayer junctional fluorescence intensities for E-cadherin, phosphorylated Myosin Light Chain (pMyosin), Desmoplakin, and Vinculin were quantified by drawing segmented line along junctions and measuring mean gray value. pMyosin and Vinculin junctional intensities were measured using E-cadherin as a landmark in a separate channel. Interlayer E-cadherin and Desmoplakin intensities were quantified in orthogonal slices by drawing segmented lines between the two epidermal layers and measuring the mean gray value for each SEC-BEC interface. Collagen and Laminin fluorescence intensities were quantified by drawing a fixed-size rectangle region of interest starting at the edge of the fin fold, followed by the plot profile function in ImageJ. The region of interest was kept consistent within each experiment. Average intensities of Integrin β1b and BEC actin levels were measured by drawing regions of interest outlining individual epidermal cells using the polygon selection tool in ImageJ. SEC actin microridge length was quantified using ImageJ as previously described^38^. Briefly, cell outlines were traced using the polygon selection tool manually to measure apical cell area. Brightness and contrast were automatically enhanced, and surrounding area was cleared. Images were blurred with the Smoothen function three times and processed through a Laplacian morphological filter from the MorphoLibJ plugin^78^ (square element, radius = 1). Filtered images were automatically thresholded using the Triangle method and skeletonized. Microridge length was measured using the Analyze Skeleton (2D/3D) feature.

#### Cell Culture Confocal Fluorescence Images

Confocal fluorescence images were analyzed with the Fiji version of ImageJ (NIH)^77^. Confocal z-stack images were taken from the suprabasal layer to the basal layer. Suprabasal and basal keratinocytes were identified by maximum projection of selected z slices based on their z-position. Intralayer junctional fluorescence intensity for β-catenin and Desmoplakin were quantified by drawing segmented line along junctions and measuring mean gray value. Interlayer β-catenin and Desmoplakin intensity were quantified by measuring mean gray value of regions of interest in single z slices at the suprabasal-basal keratinocytes interface. Interlayer Desmoplakin plaque area was quantified by first thresholding based on intensity, followed by the analyze particles function in ImageJ. P63 intensity in suprabasal and basal layers was quantified by measuring the mean gray value of regions of interest created using intensity-thresholded nuclei in maximum projection images of each keratinocyte layer.

#### Dragon-Tail Analysis using Imaris Software

Epithelial cell migration in the time lapse movies of wild type fin fold development was analyzed using Imaris spot module with manual spot tracing function. Migration distance (Track Length) was visualized as dragon tails. The average speed, track length, and duration of tracks were exported for statistical analysis.

#### Calculation for Mean Squared Displacement (MSD)

Movement profiles, time and x,y position, for individual cells in the fin fold were used to calculate a mean squared displacement (MSD) value as performed previously^79^. MSD values are defined as follows:

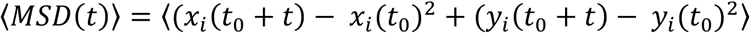

where *x*_*i*_ and *y*_*i*_ are the coordinates of the *i*th cell and *t* represents the time interval.

#### Transcriptome Analysis

Transcriptomic data were extracted from Cokus et al, 2019. In brief, distinct epithelial cell populations were FACS sorted, quality checked, and prepared with Illumina TruSeq Library. One library was used per replicate, at a sequencing depth of 21.2-45.4 million reads pass filter (PF). CutAdapt 1.8.1 was used for quality control, and reads were aligned to Ensembl release 92 Danio rerio GRCz11 using STAR 2.5.3a. Library distributions were performed with Salmon 0.10.2. Analyses were performed using DeSeq2, using a variation of Sleuth analysis described in Cokus et al 2019 Supplementary File 1.

Enrichment plots for collagen and laminin expression in different cell types were extracted from DanioCell^28^, an online database of single-cell transcriptomic data across different developmental stages. In brief, cell-type specific clusters were first identified using cell-type specific markers (*tp63* for BEC and *pdgfrb* for Mesenchymal cells). Expression patterns of different ECM proteins were then extracted in the respective clusters.

#### Force Inference Analysis

Junctional tension inference was performed as previously described in Kong et al 2019^80^. Briefly, maximum intensity projection images of fin epithelia were processed in Fiji by adjusting intensity thresholds. Segmentation was carried out using the Fiji plugin Tissue Analyzer, which provided the coordinates of cell vertices and boundaries. These data were extracted using a custom MATLAB script and analyzed with a previously published Bayesian inference algorithm to compute junctional tensions^80^. Inferred tension values for each junction were exported for downstream analysis. For statistical comparison, data from all biological replicates were pooled. A two-tailed *t*-test was used to compare inferred junctional tensions between the central and peripheral regions of the fin fold.

### Statistics and reproducibility

All statistical analyses were performed using prism version 10.4.1 (Graphpad). Measurements were taken from distinct samples. Sample size and statistical tests are specified in each figure legend. All sample size were determined using power analysis test. For all data sets, outliers were first identified and removed using the ROUT method in prism. Then, data sets were tested for Gaussian distribution using the D’Agostino-Pearson omnibus normality test (significance level = 0.05). For normally distributed datasets, unpaired t-test (two-sided) was used to test mean differences between two groups and ordinary one-way ANOVA with Tukey’s multiple comparisons test was used to test mean differences among multiple groups. For data sets that are not normally distributed, Mann-Whitney test (two-sided) was used for comparison between two groups and Kruskal-Wallis with Dunn’s multiple comparisons test was used for comparisons among multiple groups.

## Supplementary Information

**Supplementary Video 1. Synchronous Migration and Sustained Interlayer Contact of BECs and SECs in the Fin Fold during Development.**

Representative time-lapse video showing both superficial epidermal cells (SECs), labeled with *Tg(krt5:Gal4ff)^la2^*^12^*; Tg(UAS:LifeAct-GFP)^mu2^*^71^, and basal epidermal stem cells (BECs), co-labeled with *cdh1-tdTomato^xt^*^18^. Video was pseudo-colored using Imaris software, and dragon-tail tracking measured the migration distance (Track Length) of each SEC and BEC over an ∼8-hour recording period starting at 48 hours post-fertilization (hpf). Track length is displayed using a color bar scale. Individual SECs and BECs tracked in different regions of the fin fold are labeled with circles (SEC = empty circle, BEC = filled circle, Center = green, Periphery = purple).

**Supplementary Video 2. Differential Wound Healing in Peripheral vs. Central SECs**

Representative live imaging confocal time-lapse videos of *wild type*; *Tg(krt5:Gal4ff)^la2^*^12^*; Tg(UAS:LifeAct-GFP)^mu2^*^71^ for LifeAct-GFP under the SEC-specific *krt5* promoter, showing the central and peripheral regions of the fin fold at 2 dpf before and after single-cell ablation. Scale bar = 20 µm.

**Supplementary Video 3. Delayed Wound Closure in Peripheral *lama5*^+/-^ SECs Compared to *WT***

Representative live imaging confocal time-lapse videos of *Tg(krt5:Gal4ff)^la2^*^12^*; Tg(UAS:LifeAct-GFP)^mu2^*^71^ for LifeAct-GFP under the SEC-specific *krt5* promoter in the peripheral regions of *WT* vs. *lama5^+/-^* fin fold at 2 dpf before and after single-cell ablation. Scale bar = 20 µm.

**Supplementary Video 4. No Effect on Wound Closure in Central *lama5*^+/-^ SECs Compared to *WT***

Representative live imaging confocal time-lapse videos of *Tg(krt5:Gal4ff)^la2^*^12^*; Tg(UAS:LifeAct-GFP)^mu2^*^71^ for LifeAct-GFP under the SEC-specific *krt5* promoter in the center regions of *WT* vs. *lama5^+/-^* fin fold at 2 dpf before and after single-cell ablation. Scale bar = 20 µm.

**Supplementary Video 5. No Effect on Wound Closure in Peripheral *col1a1a*^+/-^ SECs Compared to WT**

Representative live imaging confocal time-lapse videos of *Tg(krt5:Gal4ff)^la2^*^12^*; Tg(UAS:LifeAct-GFP)^mu2^*^71^ for LifeAct-GFP under the SEC-specific *krt5* promoter in the periphery regions of *WT* vs. *col1a1a^+/-^* fin fold at 2 dpf before and after single-cell ablation. Scale bar = 20 µm.

**Supplementary Video 6. Delayed Wound Closure in Central *col1a1a*^+/-^ SECs Compared to WT**

Representative live imaging confocal time-lapse videos of *Tg(krt5:Gal4ff)^la2^*^12^*; Tg(UAS:LifeAct-GFP)^mu2^*^71^ for LifeAct-GFP under the SEC-specific *krt5* promoter in the center regions of *WT* vs. *col1a1a^+/-^* fin fold at 2 dpf before and after single-cell ablation. Scale bar = 20 µm.

**Supplementary Video 7. No Effect on Wound Closure in Peripheral *dspa*^+/-^ SECs Compared to WT**

Representative live imaging confocal time-lapse videos of *Tg(krt5:Gal4ff)^la2^*^12^*; Tg(UAS:LifeAct-GFP)^mu2^*^71^ for LifeAct-GFP under the SEC-specific *krt5* promoter in the periphery regions of *WT* vs. *dspa^+/-^* fin fold at 2 dpf before and after single-cell ablation. Scale bar = 20 µm.

**Supplementary Video 8. Partial Rescue of Wound Closure in Peripheral *lama5*^+/-^, *dspa*^+/-^ SECs Compared to *lama5^+/-^***

Representative live imaging confocal time-lapse videos of *Tg(krt5:Gal4ff)^la2^*^12^*; Tg(UAS:LifeAct-GFP)^mu2^*^71^ for LifeAct-GFP under the SEC-specific *krt5* promoter in the peripheral regions of *lama5^+/-^* vs. *lama5^+/-^, dspa^+/-^* fin fold at 2 dpf before and after single-cell ablation. Scale bar = 20 µm.

## Acknowledgements

We would like to thank Nicole Semanchik and Jared Hintzen for zebrafish husbandry and technical assistance with this project. We are also grateful to Robert Lalonde and Andrew Prendergast of the Yale Zebrafish Research Core for their technical service with CRISPR/Cas9 line genotype. We thank Valentina Greco, Scott Holley, and Anjelica Gonzalez at Yale University for their valuable feedback and suggestions during the development of the project. We also acknowledge Angela Andersen at Life Science Editors for editorial assistance. S.N. discloses support for this work from NHLBI R21HL165342, NHLBI P01HL169168, and AHA award 957692. K.S. discloses support from NIGMS R35GM150645.

## Author Contributions

H.M.H., L.C.B., K.S., and S.N. conceptualized and supervised the study. H.M.H. performed and analyzed the zebrafish experiments. L.C.B. performed and analyzed the 3D keratinocyte model experiments in collaboration with X.G. and M.M. J.M.B., S.L.I., and Z.W. contributed to experimental execution and data analysis. H.M.H., M.C., and S.N. wrote the manuscript with input from all authors.

---

**Extended Data Fig. 1.**
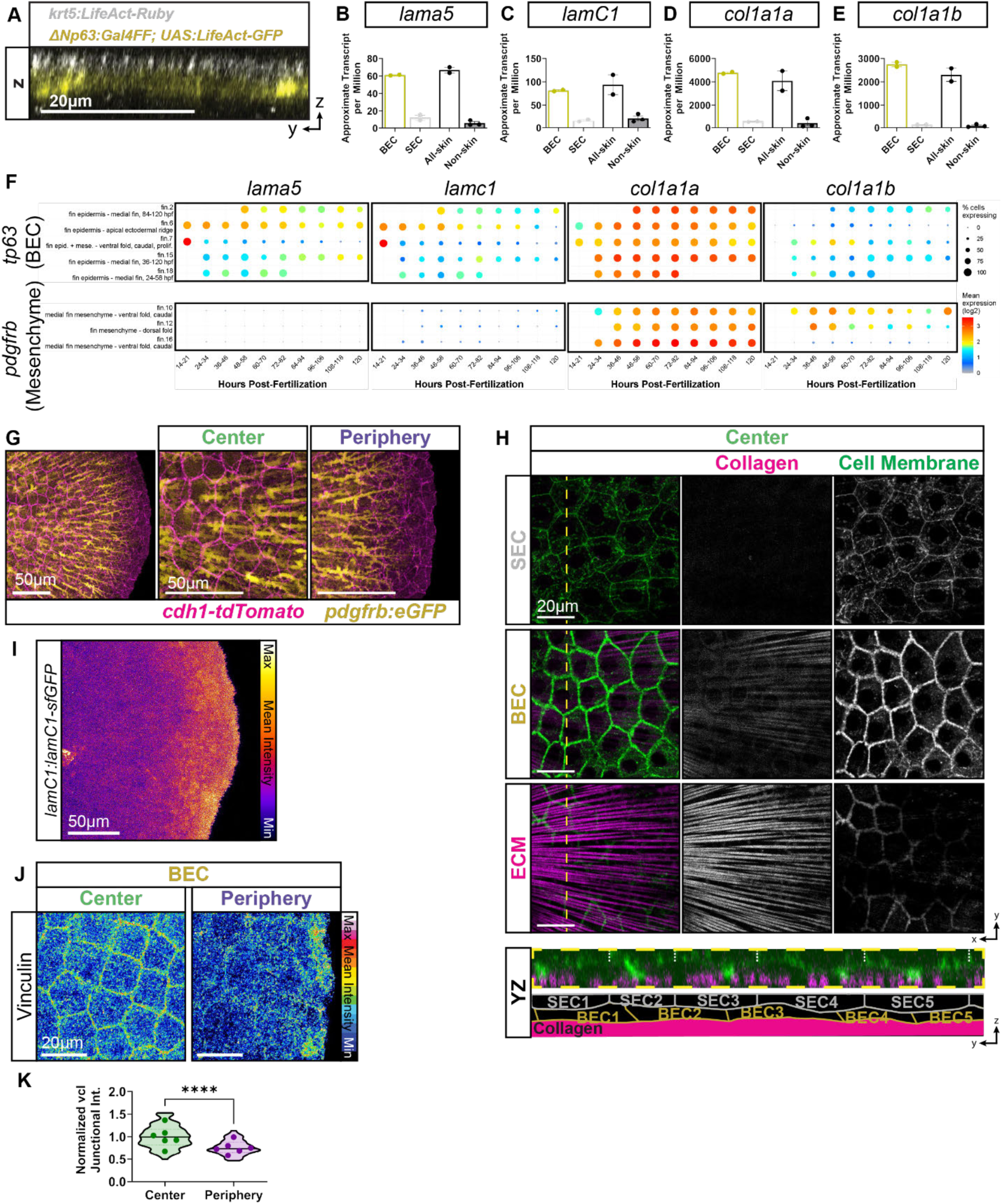
Differential ECM Composition in the Fin Fold, with Collagen Produced by Both BECs and Mesenchymal Cells in the Center and Laminin Produced by BECs in the Periphery. (**A**) Representative transverse live imaging confocal image of zebrafish embryos around 2 dpf fin fold characterized by basal epidermal stem cells (BECs, yellow) with *TgBAC(ΔNP63:Gal4ff)^la2^*^13^*; Tg(UAS:LifeAct-GFP)^mu2^*^71^ and the periderm, a layer of superficial epidermal cells (SECs, white) with *Tg(krt5:Gal4ff)^la2^*^12^*; Tg(UAS:LifeAct-GFP)^mu2^*^71^. (**B-E**) Bar plots representing the expression of the indicated genes analyzed via transcriptomic analysis of BECs and SECs sorted from skin and non-skin tissue in zebrafish embryos around 2 dpf reported in ref^21^. (**F**) Gene expression dot plot of ECM genes (*lama5, lamc1, col1a1a,* and *col1a1b*) in BECs (*tp63* enriched clusters) and mesenchymal cells (*pdgfrb* enriched clusters), reported in single-cell gene expression data from ref^28^. (**G**) Representative live imaging confocal images demonstrating localization of mesenchymal cells in different regions of the fin fold at ∼48 hpf. Mesenchymal cells visualized with *TgBAC(pdgfrb:eGFP)^ncv22Tg^,* pseudo colored in yellow and epidermal cells visualized with *cdh1-tdTomato^xt^*^18^, pseudo colored in magenta. (**H**) Representative immunofluorescence confocal z-stack of the fin fold of transgenic fish Tg(β-actin:Hras-eGFP)*^vu1^*^19^, labeling all cell membrane and stained with a Cy3-tagged collagen hybridizing peptide (labels all types of remodeled Collagen) in the central region of different epidermal layers. (Bottom) Schematic of ECM (magenta), BEC (yellow), and SEC (grey) cell membranes (green) from transverse position (yellow dashed line) (**I**) Representative live imaging confocal image of *TgBAC(lamC1:lamC1-sfGFP)p1* transgenic line showing the entire fin fold at ∼48 hpf. (**J**) Representative immunofluorescence confocal images of Vinculin in the central and peripheral regions of fin fold BECs at ∼48 hpf. Fluorescence intensity is displayed using a 16-color scale. (**K**) Quantification of fluorescence intensity of BEC junctional Vinculin across embryos, with the distribution of individual junctional measurement shown as violin plots and average values per embryo as dots for both central and peripheral regions. Intensity was normalized to the Center group (two-tailed unpaired t test on violin plot, p < 0.0001, n = 116-120 junctions, dots n = 6 embryos; 1 independent experiment).

**Extended Data Fig. 2.**
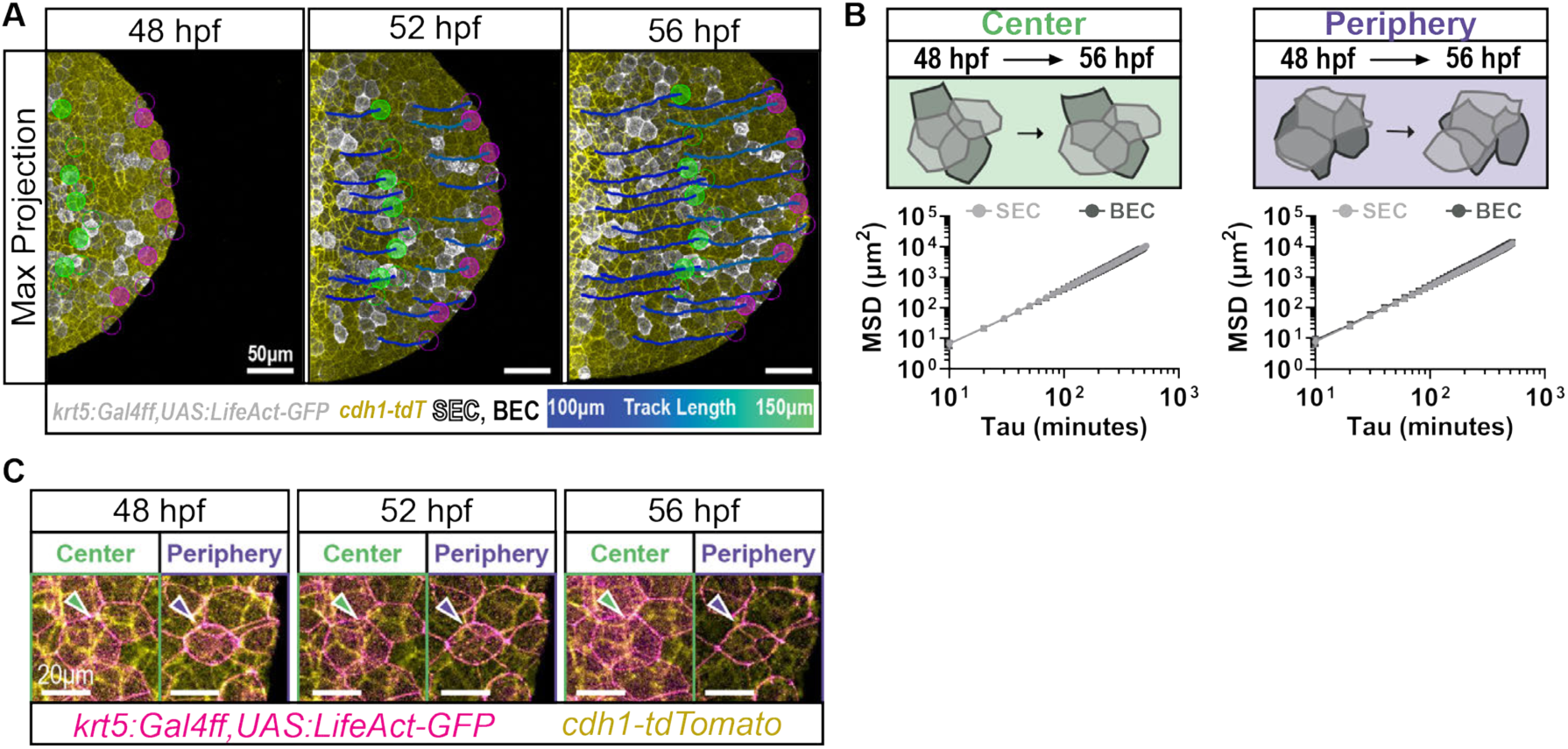
Synchronous Migration and Sustained Interlayer Contact of BECs and SECs in the Fin Fold during Development. (**A**) Representative time-lapse confocal stills showing both superficial epidermal cells (SECs), labeled with *Tg(krt5:Gal4ff)^la2^*^12^*; Tg(UAS:LifeAct-GFP)^mu2^*^71^, and basal epidermal stem cells (BECs), co-labeled with *cdh1-tdTomato^xt^*^18^. Images were pseudo-colored using Imaris software, and dragon-tail tracking measured the migration distance (Track Length) of each SEC and BEC over an ∼8-hour recording period starting at 48 hours post-fertilization (hpf). Track length is displayed using a color bar scale. Individual SECs and BECs tracked in different regions of the fin fold are labeled with circles (SEC = empty circle, BEC = filled circle, Center = green, Periphery = purple). (**B**) Schematic representation and quantification of the mean squared displacement (MSD) between SECs and the underlying BECs during fin fold development in central and peripheral regions; 3 independent experiments. (**C**) Representative time-lapse confocal stills showing superficial epidermal cells (SECs, magenta only), labeled with *Tg(krt5:Gal4ff)^la2^*^12^*; Tg(UAS:LifeAct-GFP)^mu2^*^71^, and basal epidermal stem cells (BECs, yellow without magenta), co-labeled with *cdh1-tdTomato^xt^*^18^. Over an ∼8-hour recording period starting at 48 hpf, SECs and BECs maintain their positions in both central and peripheral regions (arrowheads).

**Extended Data Fig. 3.**
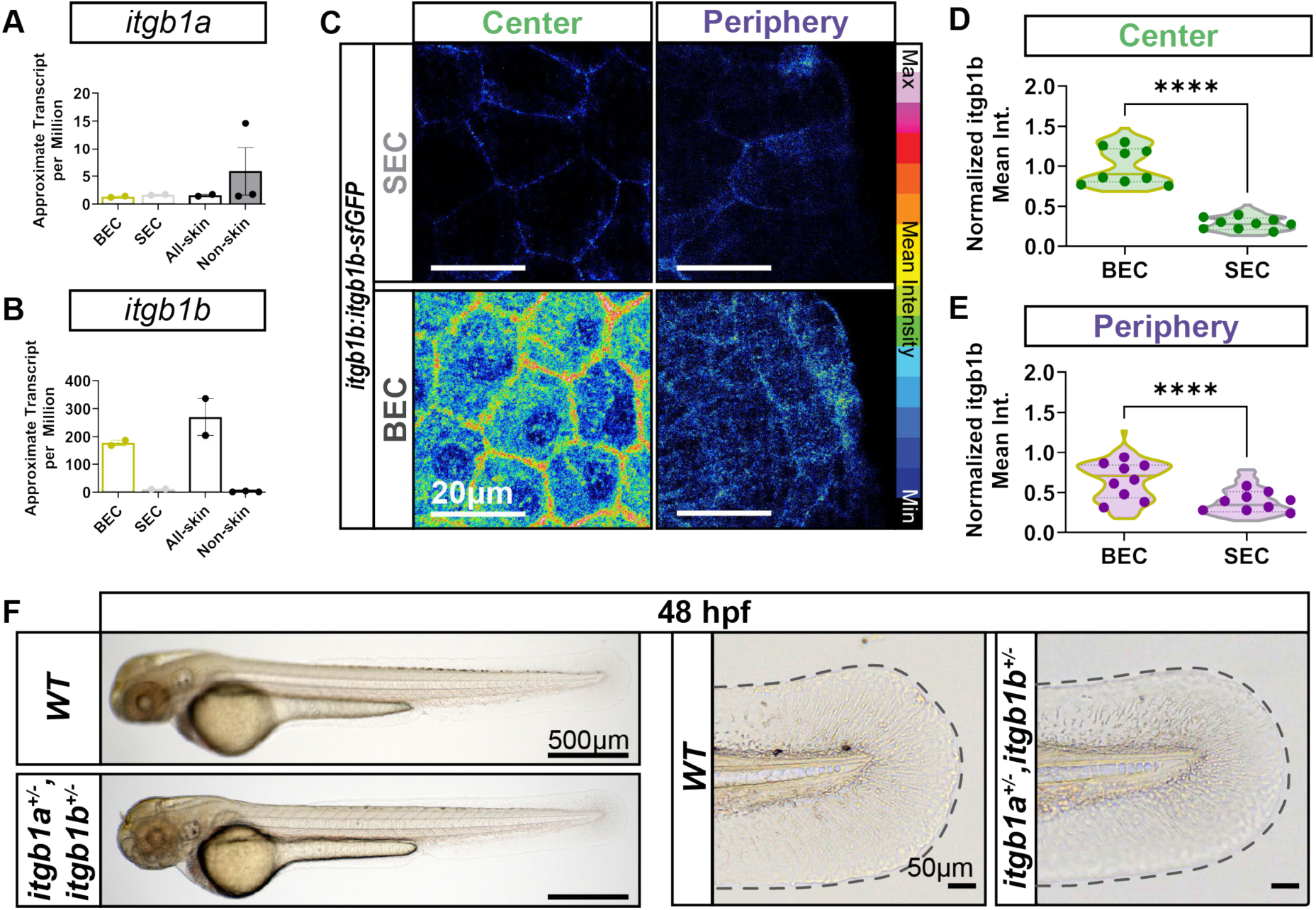
Differential Expression of *itgb1b* in BECs Across Fin Fold Regions, with Low Expression in SECs. (**A, B**) Expression levels of *itgb1a* and *itgb1b* were determined at the mRNA level using transcriptomic data reported in ref^21^ from SECs, BECs, and non-skin cells. (**C**) Representative live imaging confocal images showing *itgb1b:itgb1b-sfGFP* endogenous knock-in line of Integrin β1b fusion protein localization in SECs and BECs within the zebrafish fin fold at ∼48 hpf. Fluorescence intensity is displayed using a 16-color scale. (**D, E**) Quantification of fluorescence intensity of Integrin β1b levels per cell across embryos, with the distribution of individual cell shown as violin plots and average values per embryo as dots for both central and peripheral regions. Intensity was normalized to the BEC Center group for both graphs (two-tailed Mann-Whitney test on violin plots, p < 0.0001, n = 73-75 cells, dots n = 9 embryos; 2 independent experiments). (**F**) Representative lateral-view brightfield images of *wild type* (*WT*) and *itgb1a^+/-^, itgb1b^+/-^* mutant zebrafish embryos at ∼48 hpf. Fin fold magnification shows no gross developmental defects. Dashed black lines outline the fin fold boundary.

**Extended Data Fig. 4.**
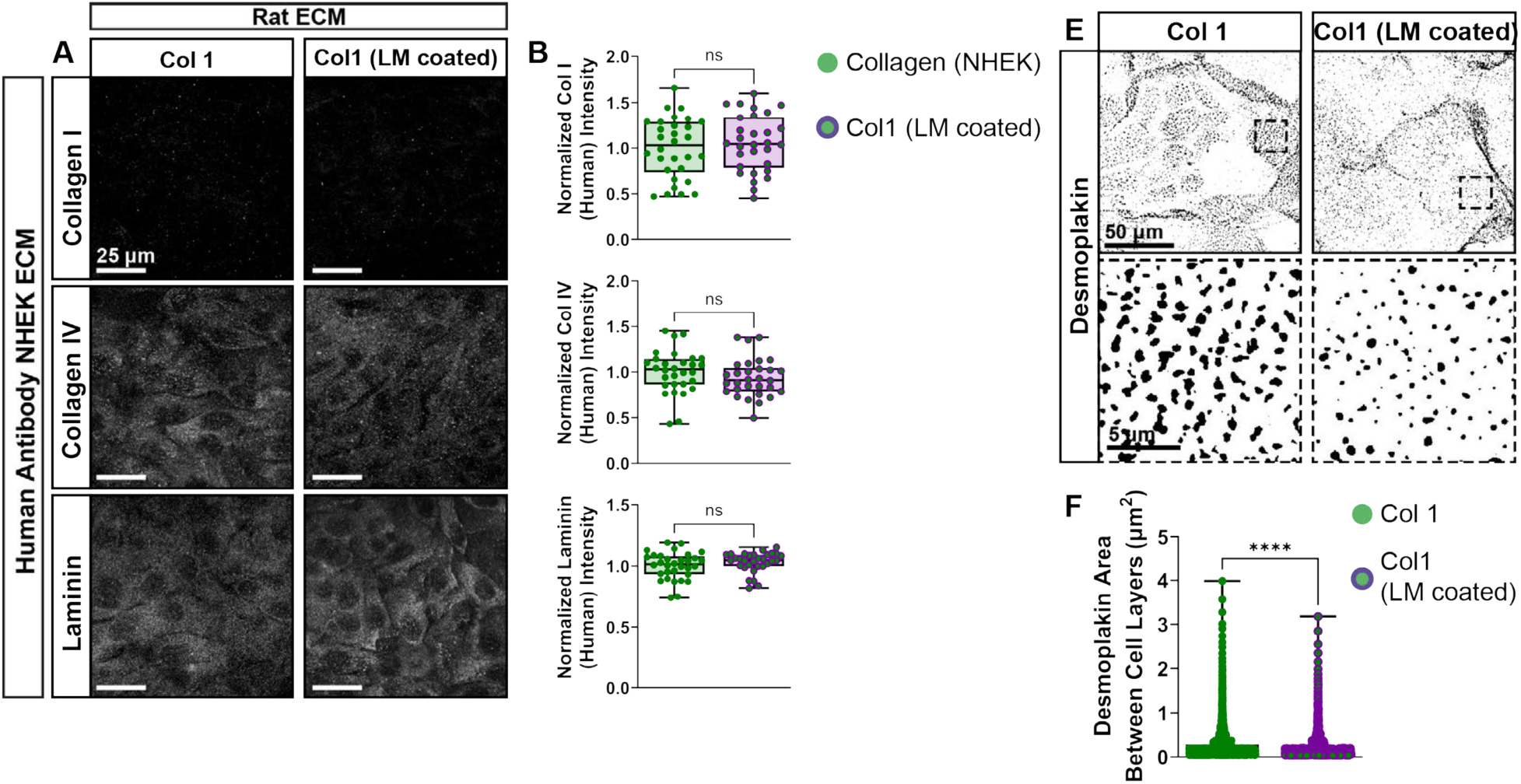
Laminin Reduced Interlayer Desmoplakin Plaque Size in Human Epidermal Keratinocytes *in vitro*. (**A**) Representative images of immunofluorescence staining specific to Human ECM proteins (rows), produced by the NHEKs when cultured on different animal-derived ECM scaffolds (columns). (**B-D**) Quantification of respective Human ECM proteins produced by NHEKs cultured on each of the gel systems (see Figure 4). (two-tailed Welch’s t test, p = 0.5393 (Collagen I), 0.2503 (Collagen IV), and 0.2227 (Laminin), n = 30-32 Field of views; 1 independent experiment). (**E**) Threshold images of interlayer Desmoplakin plaques in NHEKs cultured on each of the gel systems. (**F**) Quantification of Desmoplakin area between cell layers normalized to the collagen gel condition (Mann-Whitney test, p<0.0001, n= 936-1211 plaques; 1 independent experiment).

**Extended Data Fig. 5.**
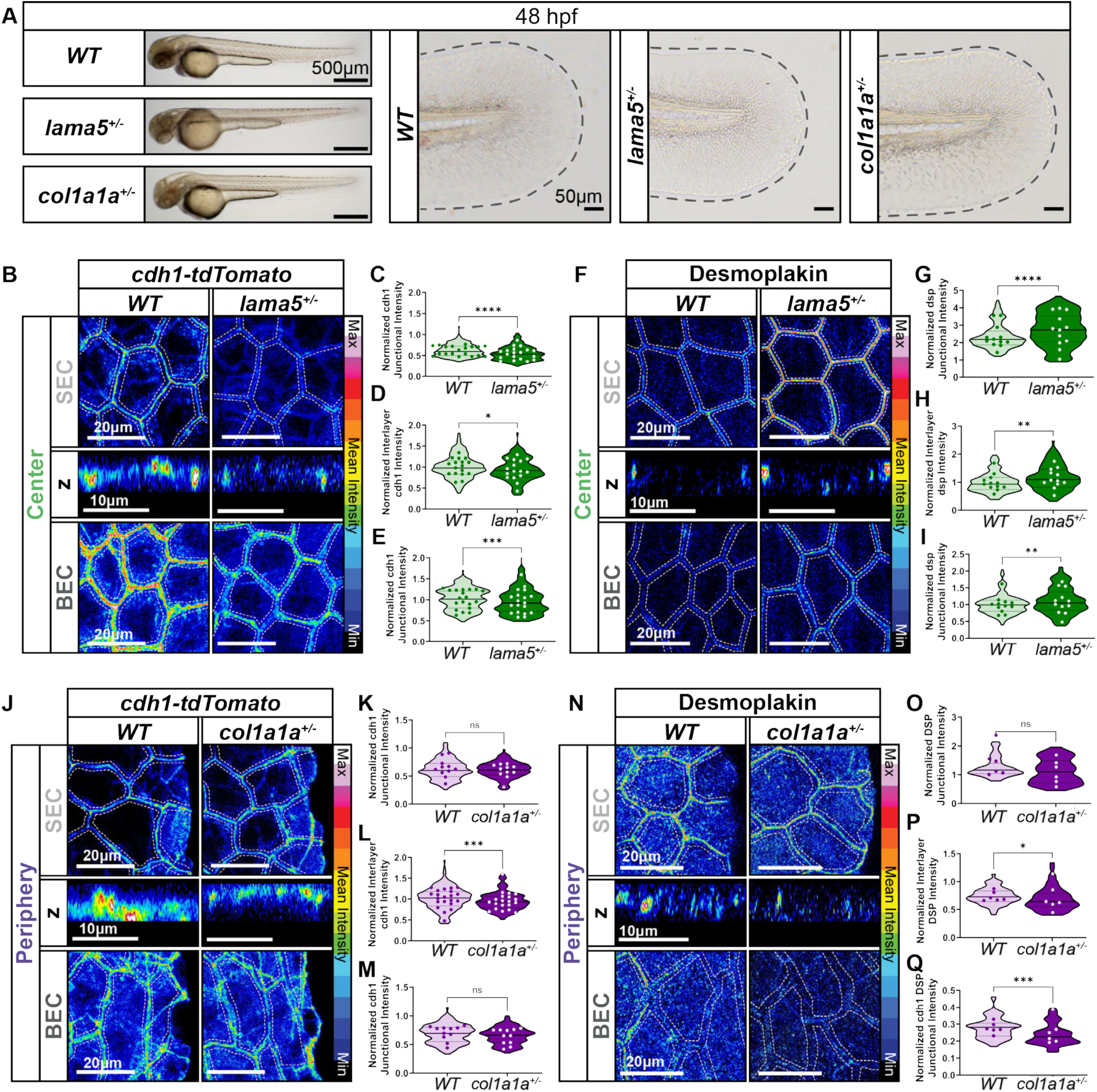
Characterization of Central BECs and SECs in *lama5* and *col1a1a* Heterozygous Zebrafish. (**A**) Representative images of *wild type* (*WT*), *lama5 and col1a1a* heterozygous zebrafish embryos at ∼48 hpf. Fin fold (dashed black lines) magnification shows no gross developmental defects. (**B, F**) Representative (**B**) images of *cdh1-tdTomato^xt^*^18^ and (**F**) desmoplakin in the central region of zebrafish fin folds at ∼48 hpf. Images show SECs, Interlayer, and BECs. Z-stack images were taken from the SEC layer to the BEC layer. SECs and BECs were identified by maximum projection of selected z slices based on their z-position. Representative interlayer images are shown in orthogonal view of single z slice. Intensity displayed using 16-color scale. (**C-E, G-I**) Quantification of (**C-E**) E-cadherin and (**G-I**) desmoplakin at central SEC junctions, interlayer, and BEC junctions with individual junctional measurement distribution shown as violin plots and average values per embryo as dots. (**C-E**) For all graphs, E-cadherin intensity was normalized to the *WT* Center BEC group (two-tailed unpaired t test used for Center Interlayer comparison and two-tailed Mann-Whitney test for all other comparisons on violin plots, p <0.0001 (SEC), p = 0.0306 (Interlayer) and 0.0004 (BEC), n = 373 - 380 junctions (SEC and BEC junctions) and 160-170 cells (interlayer), dots n = 19 (*WT*) and 19 (*lama5^+/-^*) embryos; 3 independent experiments). (**G-I**) For all graphs, desmoplakin intensity was normalized to the *WT* Center BEC group (two-tailed Mann-Whitney test on violin plots, p<0.0001 (SEC), p = 0.0025 (Interlayer) and 0.0019 (BEC), n = 240 junctions (SEC and BEC junctions) and 119-120 cells (interlayer), dots n = 12 (*WT*) and 12 (*lama5^+/-^*) embryos; 2 independent experiments). (**J, N**) Representative (**J**) images of *cdh1-tdTomato^xt^*^18^ and (**N**) desmoplakin in the peripheral region of zebrafish fin folds at ∼48 hpf. Images show SECs, Interlayer, and BECs. Z-stack images were taken from the SEC layer to the BEC layer. SECs and BECs were identified by maximum projection of selected z slices based on their z-position. Representative interlayer images are shown in orthogonal view of single z slice. Intensity displayed using 16-color scale. (**K-M, O-Q**) Quantification of (**K-M**) E-cadherin and (**O-Q**) desmoplakin at peripheral SEC junctions, interlayer, and BEC junctions across *WT* and *col1a1a*^+/-^ mutant embryos, with individual junctional measurements distribution shown as violin plots and average values per embryo as dots. (**K-M**) For all graphs, E-cadherin intensity was normalized to the *WT* Center BEC group in bottom panel of Fig.5I (two-tailed Mann-Whitney test on peripheral BEC and unpaired t test on all other violin plots, p = 0.5137 (SEC), 0.0500 (BEC) and p < 0.0001 (Interlayer); n = 168-170 junctions (SEC and BEC junctions) and 110 cells (interlayer), dots n = 11 (*WT* and *col1a1a^+/-^*) embryos; 1 independent experiment). (**O-Q**) For all graphs, Desmoplakin intensity was normalized to the *WT* Center BEC group (two-tailed Mann-Whitney test on violin plots, p = 0.0508 (SEC), 0.0208 (Interlayer) and 0.0002 (BEC), n = 60-83 junctions (SEC and BEC junctions) and 60 cells (interlayer), dots n = 6 (*WT* and *col1a1a^+/-^*) embryos; 1 independent experiment).

**Extended Data Fig. 6.**
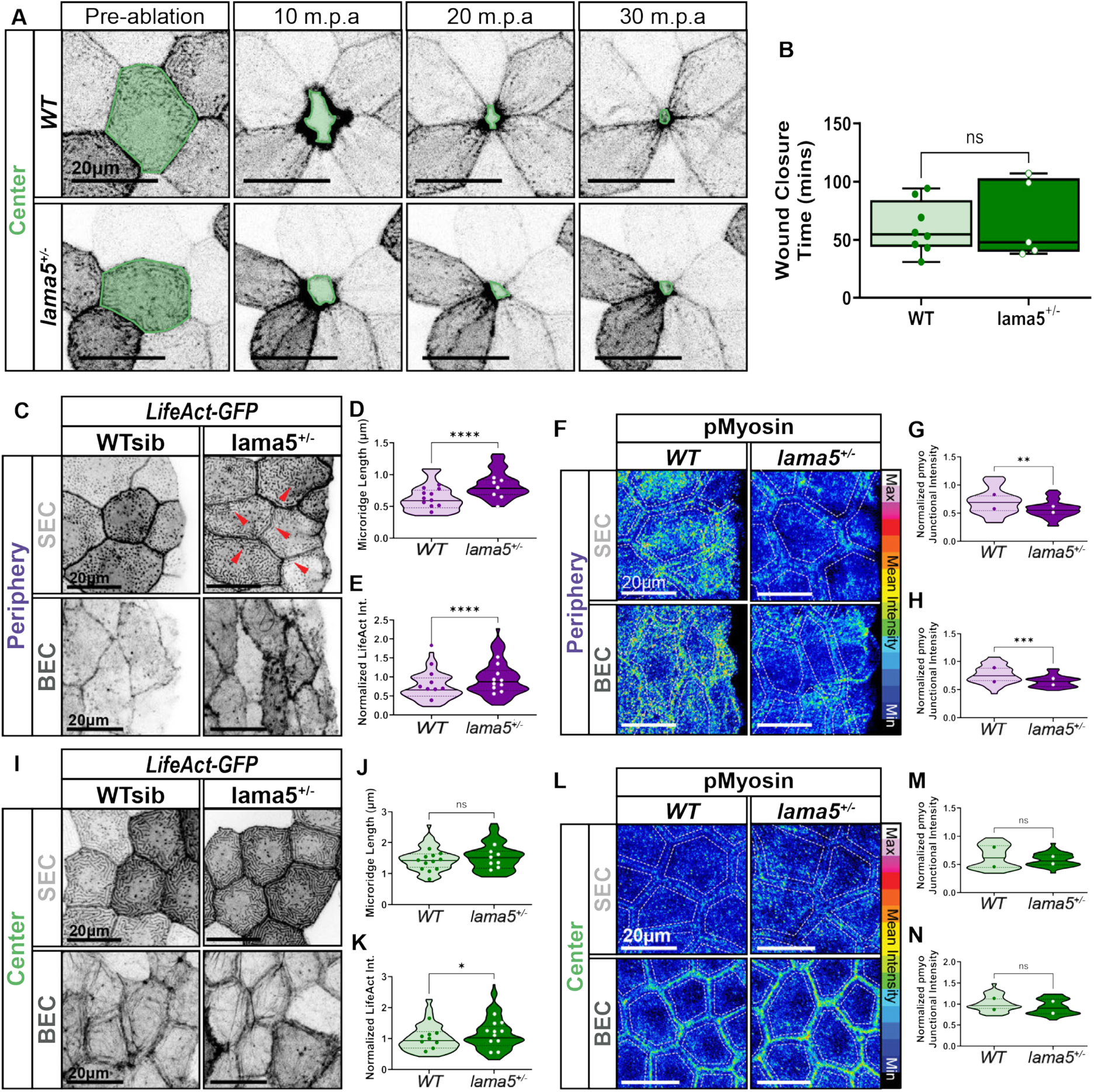
(**A**) Representative images of *Tg(krt5:Gal4ff)^la2^*^12^*; Tg(UAS:LifeAct-GFP)^mu2^*^71^ in the central regions of fin fold at 2 dpf before and after single-cell ablation. Shaded border indicates the ablated cell and subsequent wound closure. (**B)** Quantification of wound closure time (min). Box plots with min to max, each point representing indicated measurement for each ablation experiment (two-tailed Mann-Whitney test, p = 0.943279, n = 8 (*WT*) and 5 (*lama5^+/-^*) embryos; 5 independent experiments). (**C**) Representative images of *Tg(krt5:Gal4ff)^la2^*^12^*; Tg(UAS:LifeAct-GFP)^mu2^*^71^ and *TgBAC(ΔNP63:Gal4ff)^la2^*^13^*; Tg(UAS:LifeAct-GFP)^mu2^*^71^ in the peripheral regions of *WT* vs. *lama5^+/-^* fin fold at ∼48 hpf. Increased actin microridge in the *lama5^+/-^* is indicated with red arrows. (**D**) Quantifications of SEC actin microridge length with the distribution of individual cell shown as violin plots and average values per embryo as dots for peripheral region (two-tailed unpaired t test on violin plot, p<0.0001, n = 35-39 cells, dots n = 7 – 11 embryos; 2 independent experiments). (**E**) Quantification of intensity of BEC-LifeAct-GFP per cell, with the distribution of individual cell shown as violin plot and average values per embryo as dots for peripheral region. Intensity was normalized to the *WT* Center group in panel Extended Data Fig. 6K (two-tailed Mann-Whitney test on violin plot, p<0.0001, n = 110-160 cells, dots n = 9-10 embryos; 2 independent experiments). (**F**) Representative image for phosphorylated Myosin in the peripheral region at ∼48 hpf. Intensity idisplayed using 16-color scale. (**G, H**) Quantification of pMyosin at peripheral (**G**) SEC junctions and (**H**) BEC junctions, with the distribution of individual junctional measurements shown as violin plots and average values per embryo as dots (two-tailed unpaired t test on violin plots, p = 0.0021 (SEC) and 0.0001 (BEC), n = 40 junctions, dots n = 2 embryos; 1 independent experiment). For all graphs, pMyosin intensity was normalized to the *WT* Center BEC group in panel Extended Data Fig. 6N. (**I)** Representative images of *Tg(krt5:Gal4ff)^la2^*^12^*; Tg(UAS:LifeAct-GFP)^mu2^*^71^ and *TgBAC(ΔNP63:Gal4ff)^la2^*^13^*; Tg(UAS:LifeAct-GFP)^mu2^*^71^ in the center regions of *WT* vs. *lama5^+/-^* fin fold at ∼48 hpf. (**J**) Quantifications of SEC actin microridge length with the distribution of individual cell shown as violin plots and average values per embryo as dots for peripheral region (two-tailed unpaired t test on violin plot, p = 0.2237, n = 52-66 cells, dots n = 8 – 12 embryos; 2 independent experiments). (**K**) Quantification of fluorescence intensity of BEC-LifeAct-GFP per cell, with the distribution of individual cell shown as violin plot and average values per embryo as dots for the center region. Intensity was normalized to the *WT* Center group (two-tailed Mann-Whitney test on violin plot, p = 0.0222, n = 128-165 cells, dots n = 9-11 embryos; 2 independent experiments). (**L**) Representative image for phosphorylated-myosin in the center region at ∼48 hpf. Intensity displayed using 16-color scale. (**M, N**) Quantification of pMyosin at center (**M**) SEC junctions and (**N**) BEC junctions, with the distribution of individual junctional measurements shown as violin plots and average values per embryo as dots (Mann-Whitney test on violin plots, p = 0.6153 (SEC) and 0.0824 (BEC), n = 40 junctions, dots n = 2 embryos; 1 independent experiment). For all graphs, pMyosin intensity was normalized to the *WT* Center BEC group.

**Extended Data Fig. 7.**
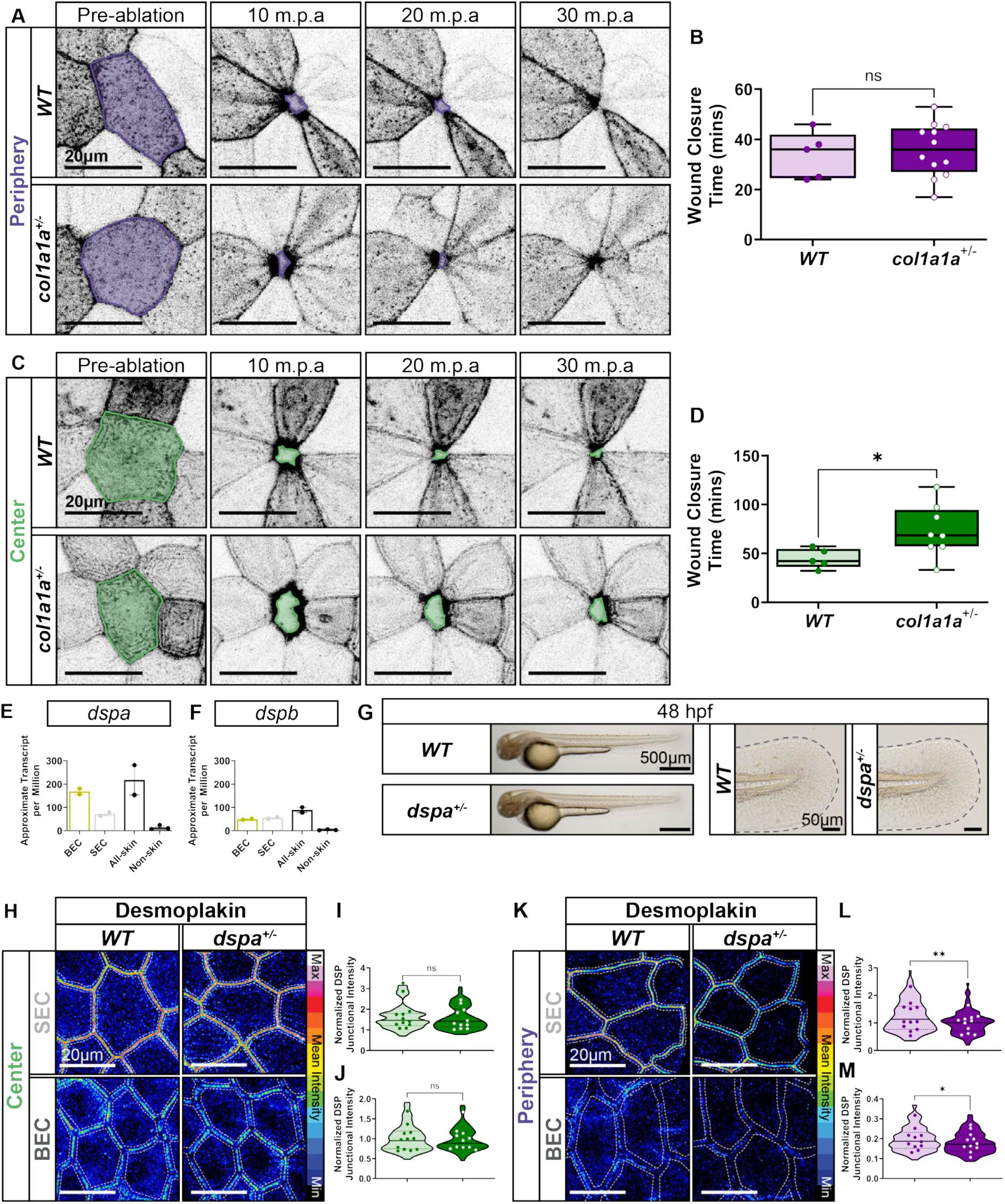
(**A**) Representative images of *Tg(krt5:Gal4ff)^la2^*^12^*; Tg(UAS:LifeAct-GFP)^mu2^*^71^ at 2 dpf before and after single-cell ablation. Shaded border indicates the ablated cell and subsequent wound closure. (**B)** Quantification of wound closure time (min),. Box plots with min to max, each point representing indicated measurement for each ablation experiment (two-tailed Mann-Whitney test, p = 0.700388, n = 5 (*WT*) and 12 (*col1a1a^+/-^*) embryos; 4 independent experiments). (**C**) Representative images of *Tg(krt5:Gal4ff)^la2^*^12^*; Tg(UAS:LifeAct-GFP)^mu2^*^71^ for LifeAct-GFP at 2 dpf before and after single-cell ablation. Shaded border indicates the ablated cell and subsequent wound closure. (**D)** Quantification of wound closure time (min),. Box plots with min to max, each point representing indicated measurement for each ablation experiment (two-tailed Mann-Whitney test, p = 0.029526, n = 5 (*WT*) and 8 (*col1a1a^+/-^*) embryos; 4 independent experiments). (**E-F**) Bar plots gene expression analyzed via transcriptomic analysis of BECs and SECs sorted from skin and non-skin tissue in zebrafish embryos around 2 dpf reported in ref^21^. (**G**) Representative images of *WT* and *dspa* heterozygous embryos at ∼48 hpf. Fin fold magnification (dashed black lines) shows no gross developmental defects.. (**H, K**) Representative staining for desmoplakin in the (**H**) central and (**K**) peripheral region of fin folds at ∼48 hpf.I Intensity displayed using 16-color scale. (**I, J**) Quantification desmoplakin at central (**I**) SEC junctions and (**J**) BEC junctions, with the individual junctional measurements distribution shown as violin plots and average values per embryo as dots (Mann-Whitney test on violin plots, p = 0.3349 (SEC) and 0.9111 (BEC), n = 108-120 junctions, dots n = 12 (*WT*) and 11 (*dspa^+/-^*) embryos). For all graphs, desmoplakin intensity was normalized to the *WT* Center BEC group in panel Extended Data Fig. 7J. (**L, M**) Quantification desmoplakin at peripheral (**L**) SEC junctions and (**M**) BEC junctions across *WT* and *dspa*^+/-^ mutant embryos, with individual junctional measurement distribution shown as violin plots and average values per embryo as dots (Mann-Whitney test on violin plots, p = 0.0050 (SEC) and 0.0283 (BEC), n = 107-120 junctions, dots n = 12 (*WT*) and 11 (*dspa^+/-^*) embryos; 1 independent experiment). For all graphs, desmoplakin intensity was normalized to the *WT* Center BEC group in panel Extended Data Fig. 7J.

## Notes

### Competing Interest Statement

The authors have declared no competing interest.

